# Diverse quorum sensing systems regulate microbial communication and biogeochemical processes in deep-sea cold seeps

**DOI:** 10.1101/2024.11.15.623595

**Authors:** Jiaxue Peng, Xinyue Liu, Jieni Wang, Nan Meng, Runlin Cai, Yongyi Peng, Yingchun Han, Jing Liao, Chengcheng Li, Maxim Rubin-Blum, Qiao Ma, Xiyang Dong

## Abstract

Quorum sensing is a fundamental chemical communication mechanism that enables microorganisms to coordinate behavior and adapt to environmental conditions. In deep-sea cold seep ecosystems, where diverse microbial communities thrive, quorum sensing is likely a key mechanism driving microbial interactions. However, the distribution, mechanisms, and ecological roles of quorum sensing in cold seeps remain poorly understood. To address this, we analyzed 173 metagenomes, 33 metatranscriptomes, and 18 metabolomes from 17 global cold seep sites. We identified 299,355 quorum sensing genes from the seep non-redundant gene catalog, representing 34 gene types across six quorum sensing systems, with distribution patterns influenced by sediment depth and seep type. These quorum sensing genes were present in 3,576 metagenome-assembled genomes from 12 archaeal and 108 bacterial phyla, revealing a complex network of intraspecies and interspecies communication. Microbial groups involved in key metabolic processes, such as sulfate-reducing bacteria, anaerobic methanotrophic archaea, diazotrophs and organohalide reducers, were extensively regulated by quorum sensing, influencing biogeochemical cycles of carbon, nitrogen, and sulfur. Phylogenetic analysis and protein domain identification highlighted the functional roles of key quorum sensing-related proteins (e.g., PDEs, CahR, RpfC/G and LuxR) in modulating microbial behaviors, such as motility and chemotaxis. Heterologous expression and metabolomic profiling further confirmed the activity of representative LuxI-R pairs and identified inhibitors of quorum sensing in cold seep sediments. Overall, these findings highlight complexity and significance of quorum sensing in microbial interactions, ecological adaptation, and biogeochemical cycling within cold seep ecosystems, advancing our understanding of microbial communication in the deep biosphere.

## Introduction

The deep-sea environment, characterized by extreme conditions such as high hydrostatic pressures, low temperatures and perpetual darkness, presents unique survival challenges for its microbial inhabitants. Despite these challenges, over 75% of marine microbial biomass thrives in the deep sea^1^. To adapt and survive in such extreme environments, microorganisms have evolved specific survival mechanisms^2^. One key mechanism is quorum sensing, a sophisticated form of cell-to-cell communication mechanism mediated by autoinducers, known as microbial languages^3^. Once cell density reaches a threshold, autoinducers are synthesized, released, accumulated, and subsequently detected through receptor-mediated recognition, facilitating the expression of multiple downstream genes and ultimately coordinating collective behaviors across the microbial community^4,5^. Collective behaviors orchestrated by quorum sensing include biofilm formation, cell competency, motility, horizontal gene transfer and symbiotic relationships, all of which are critical for the survival of deep-sea microorganisms and the maintenance of ecological functions^6–8^. Therefore, quorum sensing may exert a substantial influence on the environmental adaptability and ecological balance of deep-sea microbial communities, thereby bolstering the stability and functionality of deep-sea ecosystems.

Researchers have identified diverse quorum sensing systems and associated genes across Gram-negative and Gram-positive bacteria. The best-characterized quorum sensing system is driven by acylated homoserine lactones (AHLs)^9^, typically relying on the LuxI-R circuit to regulate cellular cooperation and communication^10^. Another important quorum sensing system, autoinducer-2 (AI-2), consists of a series of 4,5-dihydroxy-2,3-pentanedione (DPD) derivatives, generally biosynthesized from S-ribosylhomocysteine (SRH) via the enzyme LuxS, functioning as a dedicated signaling molecule for intraspecies and interspecies communication among prokaryotic organisms^11^. Other quorum-sensing systems, such as diffusible signal factors (DSF)^12^, *Pseudomonas* quinolone signals (PQS)^13^, bis-(3’-5’)-cyclic diguanylic acid (c-di-GMP)^14^, and alpha-hydroxyketones (AHKs)^15^, also facilitate intercellular communication across various microbial taxa^16^. Together, these quorum sensing systems shape microbial interactions and biogeography in natural habitats^17^, highlighting the complexity and diversity of microbial communication mechanisms.

Advancements in quorum sensing research have provided valuable insights into its contributions to marine ecosystems^18^, including regulating substance degradation, facilitating symbiosis and driving biogeochemical cycles^2^. For instance, in marine environments, quorum sensing play a role in coordinating hydrolase enzyme expression in bacteria, thereby enhancing the degradation of various polymers and influencing the cycling of carbon, phosphorus, and nitrogen^19^. In marine sulfur-oxidizing bacteria, quorum sensing regulates the synthesis of enzymes involved in the conversion of hydrogen sulfide to sulfate^2,20^. Similarly, in species like *Vibrio* and *Marinobacter*, quorum sensing governs the expression of nitrogen cycle-related genes, such as those involved in nitrogen fixation^21^, as well as genes related to hydrocarbon degradation, thereby promoting both nitrogen and carbon cycling^18,22^. In the sponge symbiont *Ruegeria* sp. KLH11, the SsaRI system provides quorum sensing-dependent control of flagellar motility functioning through the CtrA master regulator^23^. Despite extensive research on quorum sensing in coastal and shallow marine environments, the prevalence and specific mechanisms of quorum sensing in deep-sea ecosystems, particularly within cold seeps, are rarely investigated.

Cold seeps, as one of typical deep-sea environments, are located on the seafloor of continental margins with unique geological structures. These areas are formed by fluids rich in hydrocarbon gases (primarily methane), which promote the growth, specialization, and adaptation of microbial communities^24–26^. Cold seeps harbor unique and diverse microbial communities with complex microbial interactions^27^. Most importantly, the symbiosis between anaerobic methane-oxidizing archaea (ANME) and sulfate-reducing bacteria (SRB) couples anaerobic methane oxidation (AOM) with sulfate reduction, substantially contributes to carbon and sulfur cycling in cold seep ecosystems^28^. Quorum sensing is likely involved in mediating microbial interactions in cold seeps, serving as a survival strategy under extreme conditions^29,30^. However, the role of quorum sensing in microbial adaptation mechanisms and its impact on biogeochemical cycling in cold seep ecosystems remain understudied, leaving a gap in our understanding of microbial communication processes in deep-sea environments.

Here, we aimed to explore the distribution, interaction mechanisms, and ecological functions of quorum sensing in deep-sea cold seeps through a comprehensive analysis encompassing 173 metagenomes, 33 metatranscriptomes, and 18 metabolomes **(Supplementary Fig. 1**). Our findings suggest a rich diversity of quorum sensing systems in cold seep ecosystems, with a particular emphasis on the role of AI-2 signaling. These systems likely influence not only microbial cooperation and metabolic regulation but also broader biogeochemical processes. This research enhances our understanding of the ecological role of quorum sensing in the deep biosphere and offers new insights into how microbial metabolic functions and survival strategies are shaped in extreme environments.

## Results and discussion

### Cold seeps harbor diverse, actively expressed and depth-stratified quorum sensing genes

Using the quorum sensing-related proteins (QSP) database^31^ and a stringent identification workflow (**Supplementary Table 1**), we identified a total of 299,355 quorum sensing genes, representing 2.03% of the 147 million non-redundant genes in cold seeps^32^ (**Supplementary Table 2**). These genes were categorized into six quorum sensing systems based on their signaling molecules^33^ (**Fig. 1a**): (i) AI-2 (n = 137,067), (ii) DSF (n = 86,389), (iii) AHLs (n = 41,598), (iv) c-di-GMP (n = 30,584), (v) PQS (n = 2,977), and (vi) AHKs (n = 740). Functionally, these quorum sensing genes were grouped into four categories: transport (58.63%), degradation (27.71%), synthesis (12.20%) and acceptation (1.46%).

**Figure 1.**
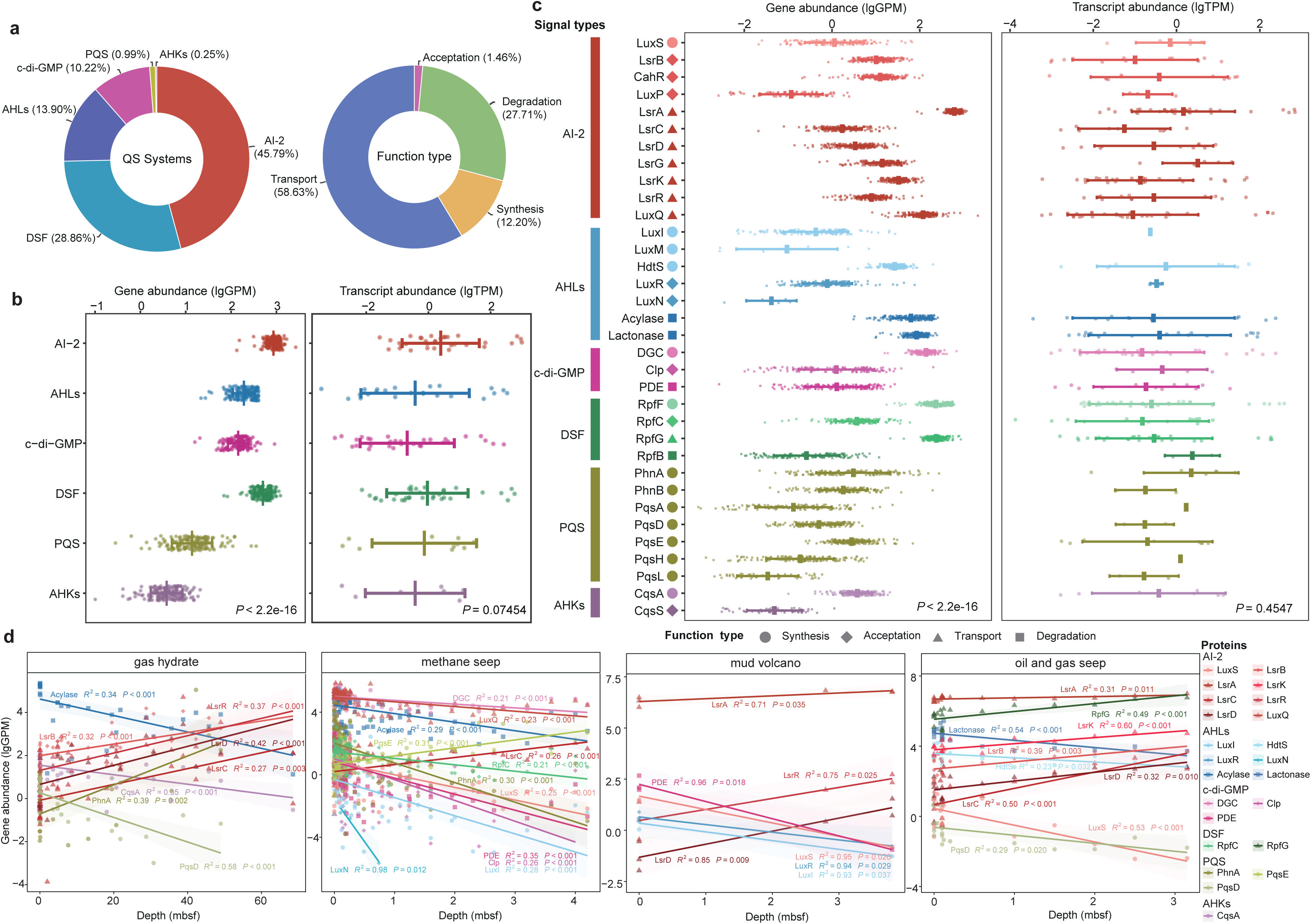
Relative abundance patterns of quorum sensing genes in cold seep sediments. **(a)** Relative proportions of 299,355 quorum sensing genes across six quorum sensing systems (left chart) and their functional roles (right chart). **(b)** Gene abundance (lg-transformed genes per million, lgGPM) and transcript abundance (lg-transformed transcripts per million, lgTPM) of six quorum sensing systems across different sediment samples. **(c)** Gene abundance (lgGPM) and transcript abundance (lgTPM) of 34 types of quorum sensing genes across various sediment samples. Different colors and shapes denote different quorum sensing systems and functions. **(d)** Relationship between quorum sensing gene abundance and sediment depth across different cold seep types. Each point represents the average gene abundance for a sample, with linear regression lines and *R²* values indicating the strength and direction of correlations. Detailed statistical data are provided in **Supplementary Tables 3-7**.

The average gene abundance of quorum sensing systems in cold seep sediments ranged from 4.98 to 912.60 GPM (genes per million), and their transcriptional abundance ranged from 4.16 to 101.59 TPM (transcripts per million) (**Fig. 1b and Supplementary Table 3**), indicating active chemical signaling regulation. Among the various types of quorum sensing genes (**Fig. 1c and Supplementary Table 4**), *lsrA*, a gene involved in AI-2 transport, exhibited the highest abundance (648.25 GPM), followed by the DSF synthetase gene *rpfF* (273.95 GPM), DSF transporter gene *rpfG* (264.56 GPM), AI-2 transporter gene *luxQ* (155.40 GPM), and c-di-GMP synthetase DGC gene (153.88 GPM). The high abundance of AI-2, DSF, AHLs, and c-di-GMP-related genes suggests that these quorum sensing signals are key mediators of microbial communication within cold seep ecosystems. Statistical analyses revealed significant differences in relative abundances of various quorum sensing systems and genes across cold seep sediments (Kruskal-Wallis, *P* < 0.01; **Fig. 1b-c and Supplementary Tables 5-6**). These differences likely reflect microbial adaptation to the cold seep environment via quorum sensing, resulting in alterations to quorum sensing systems, receptors, and downstream signaling pathways^34,35^. Additionally, genes involved in quorum quenching, such as AHL-acylase gene (94.98 GPM) and AHL-lactonase gene (88.91 GPM), were also abundant in cold seep sediments (**Supplementary Table 4**). The relatively high abundance (with a peak of 648.25 GPM) and expression levels (with a peak of 71.08 TPM) of quorum sensing and quorum quenching genes underscore that signal modulation and disruption processes collectively contribute to the regulatory dynamics of cold seep ecosystems^36^.

The distribution of quorum sensing systems in cold seeps exhibited depth-dependent trends and varied across different cold seep types (**Fig. 1d**). When samples were categorized by sediment depth (surface: < 2 meters below seafloor, mbsf; middle: 2-10 mbsf; deep: > 10 mbsf), significant differences were observed in the abundance of quorum sensing genes across depth layers (Kruskal-Wallis, *P* = 0.0043; **Supplementary Fig. 2 and Supplementary Table 7**). Notably, the relative abundance of quorum sensing signal molecule synthase and receptor genes, including *clp*, *cqsA*, *hdtS,* and *rpfC*, decreased significantly with sediment depth (*P* < 0.05). Conversely, the abundance of most transporter genes increased with sediment depth (**Fig. 1d**). This pattern may be explained by lower microbial densities and limited energy sources in deeper sediments^37^, which reduce non-essential microbial interactions and lead to decreased energy investment in signal molecule synthesis^38^. Instead, microorganisms may rely more on external signaling molecules, resulting in a reduced abundance of synthase genes and an increased abundance of transporter genes in deeper sediments^4^. Significant differences in quorum sensing systems were also observed among the five cold seep types, apart from AHLs (**Supplementary Fig. 3**). The depth distribution patterns of most quorum sensing genes varied across different cold seep types, except for the AI-2 synthase gene *luxS*, which showed a marked decline in abundance as depth increased across all seep types as depth increased (**Fig. 1d**). This suggests that the characteristic differences of each cold seep type may influence the distribution of microbial quorum sensing genes.

### Quorum sensing systems are prevalent across archaea and bacteria in cold seeps

Out of 3,813 cold seep species-level representative metagenome-assembled genomes (MAGs)^32^, 3,576 MAGs (93.78%), including 3,076 bacterial MAGs (96.91%) and 500 archaeal MAGs (78.25%), contained quorum sensing genes (**Fig. 2a and Supplementary Table 8**). These MAGs harbored a total of 32,500 quorum sensing genes spanning six quorum sensing systems and covered 34 gene types. AI-2, known for mediating both intraspecies and interspecies communication^39^, was the most widely distributed quorum sensing system, found in 83.79% of MAGs. Following it were DSF (73.25%), c-di-GMP (55.97%), AHLs (44.61%), PQS (10.78%), and AHKs (1.36%). Bacteria mainly encoded genes related to AI-2, DSF, and c-di-GMP, while archaea favored AI-2, DSF, AHLs, with limited use of c-di-GMP (only 5.63% of signal-regulated archaeal MAGs contained c-di-GMP-related genes). AHKs were exclusively found in bacterial MAGs (**Fig. 2b-c**). These findings highlight differences in quorum sensing systems between bacteria and archaea^40^.

**Figure 2.**
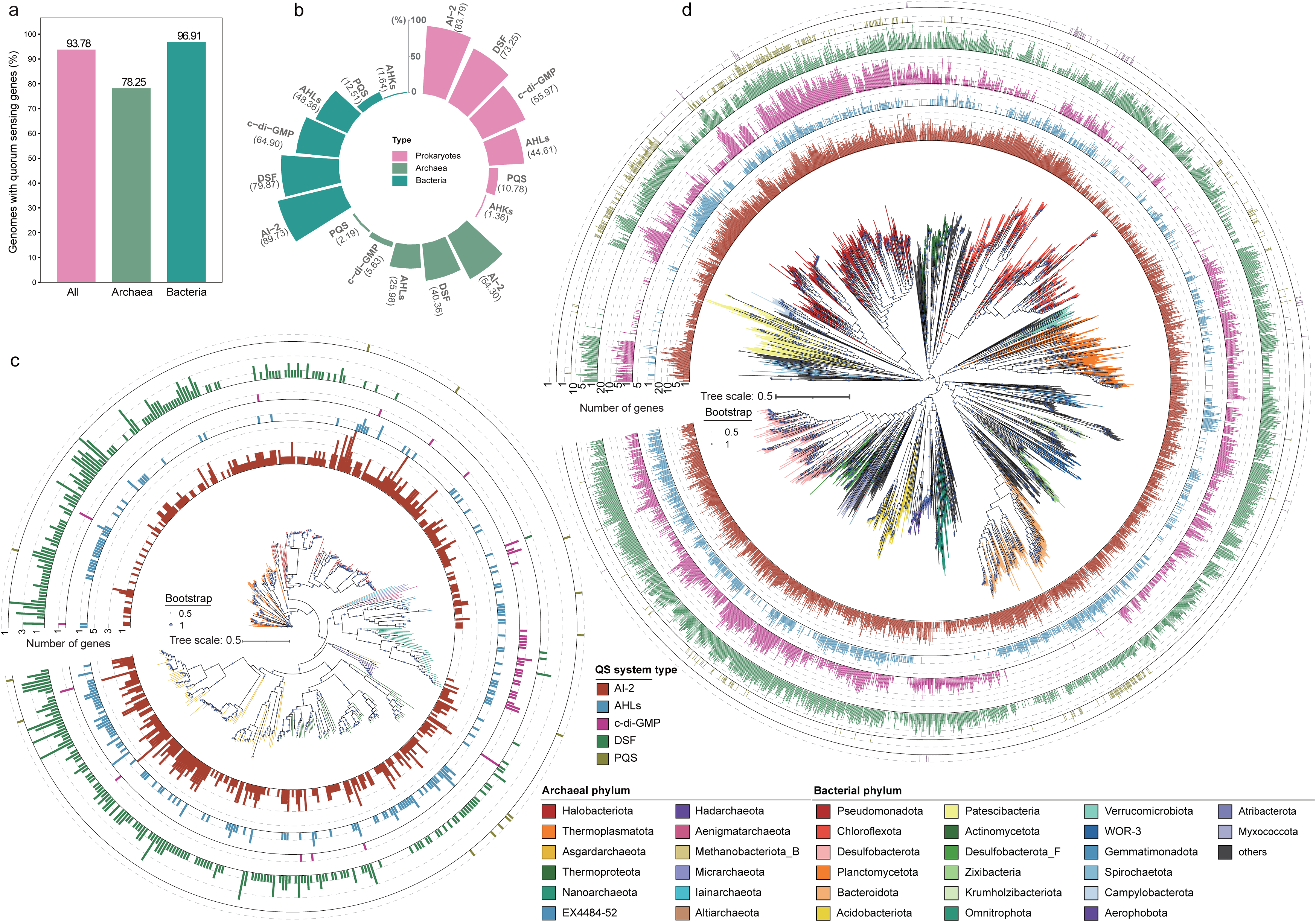
Distribution of quorum sensing genes in cold seep microorganisms. **(a)** Percentage of prokaryotic, archaeal and bacterial MAGs containing quorum sensing genes. **(b)** Proportional distribution of six quorum sensing systems among prokaryotes, archaea and bacteria. Phylogenetic trees of archaeal **(c)** and bacterial **(d)** MAGs containing quorum sensing genes. The presence of quorum sensing genes corresponding to six quorum sensing systems is indicated by colored circles on concentric rings around the tree. The outermost bar chart shows the number of quorum sensing genes related to each quorum sensing system per MAG. Detailed data are provided in **Supplementary Table 8.**

We identified quorum sensing genes in 12 archaeal phyla and 108 bacterial phyla (**Supplementary Fig. 4 and Supplementary Table 8**). In archaea, quorum sensing genes were primarily found in the phyla *Halobacteriota* (n = 117), *Thermoplasmatota* (n = 107), *Asgardarchaeota* (n = 103), *Thermoproteota* (n = 94) and *Nanoarchaeota* (n = 31). Within the *Halobacteriota*, AI-2 signaling was the most common, while most *Nanoarchaeota* lacked genes for multiple quorum sensing systems (**Fig. 2c**). Among bacteria, the majority of quorum sensing genes were found in the phyla *Pseudomonadota* (n = 471), *Chloroflexota* (n = 392), *Desulfobacterota* (n = 269), and *Planctomycetota* (n = 224) (**Fig. 2d**). Notably, *Pseudomonadota* contained a large and diverse set of quorum sensing genes (n = 8,038), with c-di-GMP-related genes being the most abundant (n = 3,301). PQS-related genes were also rich in *Pseudomonadota* but rare in other phyla, suggesting that PQS may play a key role in facilitating intraspecies communication within this group^41^. *Chloroflexota* mainly encoded genes related to AI-2 (n = 1,347) and DSF (n = 948) systems. These findings suggest that different microbial phyla in cold seeps prioritize distinct quorum sensing systems, allowing for the establishment of complex microbial interactions and specialized ecological functions^2^.

Comparing to other ecosystems, cold seep microorganisms prefer to different quorum sensing patterns. In cold seep environments, predominant AHL-active taxa include *Desulfobacterales*, *Rhodobacterales*, *Bacteroidales*, and *Flavobacteriales* (**Supplementary Table 8**), while *Rhodobacterales*, *Hyphomicrobiales*, *Enterobacterales*, *Sphingomonadales*, and *Bacillales* dominate in hydrothermal vent vent ecosystems^42^. This highlights the role of quorum sensing in modulating microbial community dynamics. Compared to rumen microorganisms (AI-2: 69.32%, DSF: 9.79%, AHKs: 1.53%)^39^, cold seep microorganisms exhibited a higher prevalence of quorum sensing genes (**Fig. 2b**), suggesting that cold seeps may act as hotspots for quorum sensing, driven by microbial adaptation to environmental pressures and resource demands. The further emphasizes how environmental factors, such as temperature and chemical composition, shape microbial community structure and signaling mechanisms.

In addition to bacterial and archaeal MAGs, we also identified 84 quorum sensing genes within 76 viral genomes (**Supplementary Table 9**). Viral quorum sensing genes were often located adjacent to genes encoding IstB-like ATP-binding proteins and anti-repressor proteins (**Supplementary Fig. 5**). These proteins are associated with functions such as stimulation of reactions driven by transposases and integrases, as well as interference with repression mechanisms^43–45^. The viral genomes were primarily associated with bacterial hosts, such as *Pseudomonas* and *Desulfobacterales* (**Supplementary Table 9**). These gene associations suggest that viruses may use quorum sensing to regulate infection and replication, potentially modulating their behavior in response to the host’s physiological state and thereby influencing the dynamics of host-virus interactions.

### Multiple quorum sensing systems mediate extensive intraspecies and interspecies communications within cold seep microbial communities

Spearman’s correlation analyses revealed that 21.75% of quorum sensing gene pairs exhibited significantly strong correlations (|*R*| ≥ 0.5, *P* < 0.05), and 96.72% of them were positive (*R* ≥ 0.5) (**Fig. 3a and Supplementary Table 10**), indicating that these quorum sensing genes are more likely to function cooperatively or co-occur in microbial communities. Strongly positive correlations were observed within AI-2, AHLs, PQS, and c-di-GMP systems. Particularly, certain signal molecule synthase and receptor genes showed strong positive correlations, such as *luxS* and *luxP* in the AI-2 system (*R* = 0.72, *P* < 0.001), and *luxI* and *luxR* in the AHLs system (*R* = 0.75, *P* < 0.001). Genes involved in the synthesis and degradation of the same signaling molecule, such as *luxI* and AHL-acylase gene (*R* = 0.65*, P* < 0.001), DGC and PDE genes (*R* = 0.60, *P* < 0.001), as well as *rpfF* and *rpfB* (*R* = 0.55, *P* < 0.001), also exhibited significant positive correlations (**Fig. 3a and Supplementary Table 10**). These results indicate that quorum sensing and quorum quenching mechanisms are tightly coordinated among cold seep microorganisms^33^. Furthermore, significantly positive correlations were observed between genes from different quorum sensing systems, such as *phnA* (PQS system) and *clp* (c-di-GMP system) (*R* = 0.84, *P* < 0.001), DGC gene (c-di-GMP system) and *luxQ* (AI-2 system) (*R* = 0.80, *P* < 0.001), and *rpfF* (DSF system) and AHL-acylase gene (AHLs system) (*R* = 0.77, *P* < 0.001). These findings suggest that multiple quorum sensing systems are co-regulate microbial behavior in cold seep ecosystems^33,46^. Additionally, some gene pairs from different systems exhibited significantly negative correlations (*R* ≤ −0.5, *P* < 0.05), such as *lsrD* (AI-2 system) and PDE gene (c-di-GMP system) (*R* = −0.53, *P* < 0.001), suggesting that these microorganisms generally use these genes separately.

**Figure 3.**
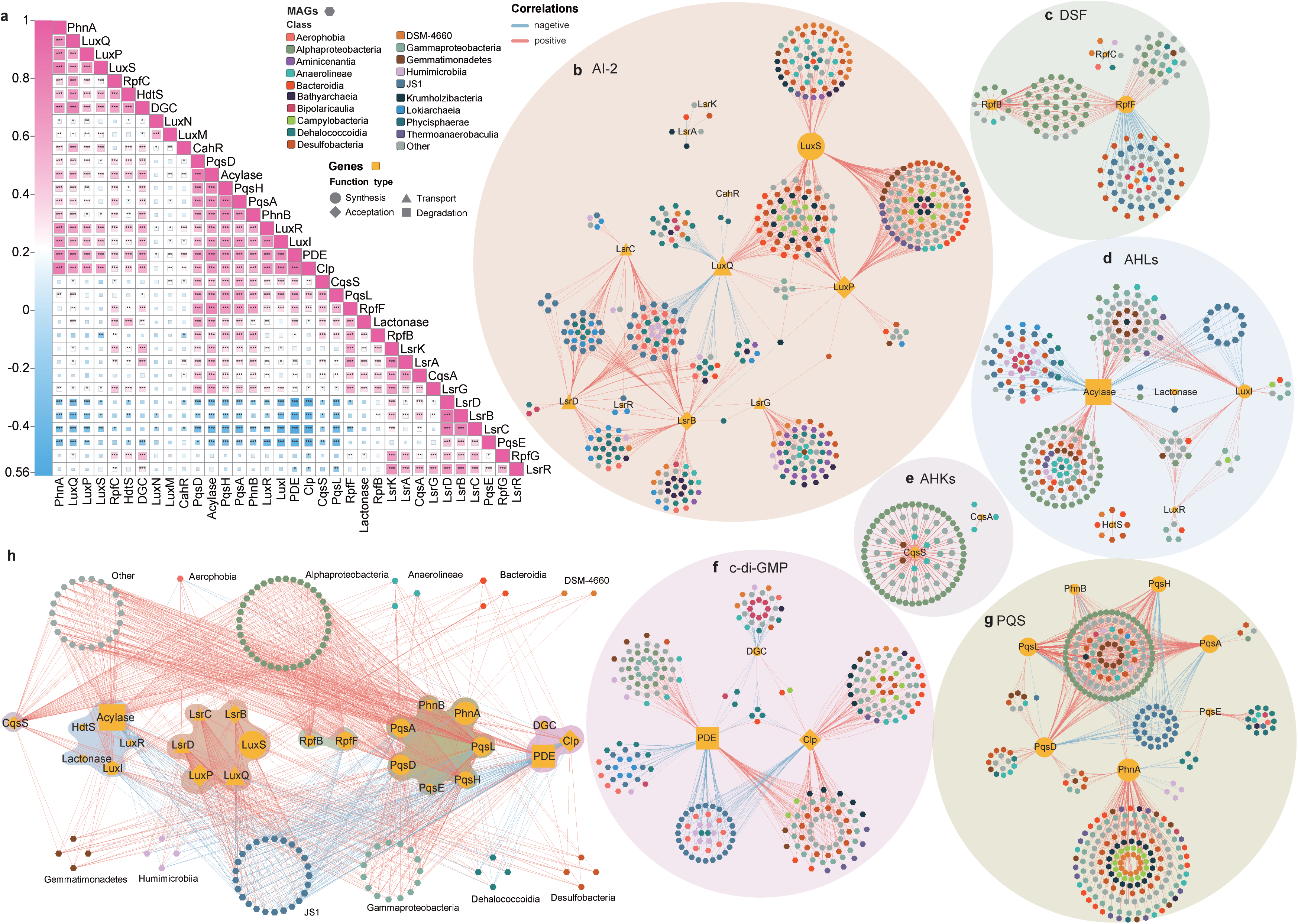
Correlation of different types of quorum sensing genes and microorganisms in cold seeps. **(a)** Spearman’s correlation coefficients among 34 types of quorum sensing genes are displayed, with positive correlations shown in shades of pink and negative correlations in shades of blue. The intensity of the color corresponds to the strength of correlations, with darker colors indicating stronger relationships. Statistical significance is denoted by asterisks: * *P* < 0.05; ** *P* < 0.01; *** *P* < 0.001. **(b)** Significant correlations between quorum sensing genes (yellow nodes) and MAGs grouped by quorum sensing systems. Nodes are colored according to microbial class, and shapes of quorum sensing gene nodes represent their functional roles. Node size reflects the degree (number of connections). Edges represent significant correlations, with red edges indicating positive correlations and blue edges indicating negative correlations. **(c)** A network based on MAGs with the highest degree of connections (degree = 8-14) and their related quorum sensing genes. Detailed statistical data are provided in **Supplementary Tables 10-13**.

Further analysis of the correlations between relative abundances of quorum sensing genes and cold seep microorganisms revealed that 696 MAGs were strongly correlated with 31 quorum sensing genes (**Fig. 3b-g and Supplementary Table 11**), with most correlations being positive. The AI-2 system exhibited the highest correlation with microbial genomes, with 495 MAGs linked to 11 AI-2-related genes (**Fig. 3b and Supplementary Table 11**). In particular, two known AI-2 response mechanisms were identified in cold seeps^47^. For the first mechanism, genes involved in AI-2 synthesis (*luxS*), transport (*luxQ*), and reception (*luxP*) formed a tightly connected group, with 63 MAGs correlated with all three genes and another 107 MAGs linked to *luxS-luxP* pair. The primary phyla represented within these groups were *Desulfobacterota* (n = 132), *Pseudomonadota* (n = 98), and *Chloroflexota* (n = 95), along with a subset of ANME (n = 10) (**Supplementary Table 12**). The strong correlation within *luxS-luxQ-luxP* cluster suggests (**Fig. 3b**) that microorganisms primarily use AI-2 for intraspecies communication. For the second mechanism, 145 MAGs were linked to the receptor gene *lsrB* and various transporters within the *lsr* operon, including *Dehalococcoidia* (n = 46)*, JS1* (n = 33), and *Lokiarchaeia* (n = 21). These microorganisms belong to different classes from those using the first mechanism, indicating that they use external AI-2, likely for interspecies communication^18^.

Many cold seep microorganisms were strongly correlated with only partial functions or individual genes within a quorum sensing system (**Fig. 3c-f**), indicating that they may not possess complete quorum sensing circuits but instead engage in “crosstalk” or “eavesdropping” for interspecies communication^48^. For example, many microorganisms, mainly *Alphaproteobacteria* (n = 42) and *Gammaproteobacteria* (n = 18), were associated with both synthase and degrading enzyme genes of DSF and AHLs systems (**Fig. 3c-d**), but fewer were linked to both synthase and receptor genes. Additionally, in the network of AHKs system (**Fig. 3e**), no MAGs were strongly correlated with both the receptor gene *cqsS* and synthase gene *cqsA*. Despite a high number of MAGs associated with c-di-GMP-related genes (average degree of genes = 132.67), 60.64% of these MAGs were correlated with only one of them (**Fig. 3f and Supplementary Table 11**). These findings suggest that cold seep microorganisms commonly use these four systems for interspecies communication. In contrast, the PQS network shows that most microorganisms, mainly *Alphaproteobacteria* (n = 48), were correlated with multiple PQS synthase genes (**Fig. 3g**). These genes tended to form clusters that facilitate the independent synthesis of PQS, which suggests that these microorganisms possess the capability to synthesize PQS^49^.

A subset of cold seep microorganisms (n = 125, 17.96%) was strongly correlated with multiple quorum sensing genes (degree = 8-14; **Fig. 3h**), suggesting they are influenced by several quorum sensing systems simultaneously. These microorganisms were primarily *Alphaproteobacteria*, *JS1*, and *Gammaproteobacteria* (**Supplementary Table 13**)*. Alphaproteobacteria* displayed strong correlations with five quorum sensing systems other than AI-2, while *JS1* primarily relied on the AI-2 system. *Gammaproteobacteria* were positively correlated with all six quorum sensing systems (**Fig. 3h**). Many of these microorganisms served as hub nodes in the microbial co-occurrence network, being highly interconnected with other linages (n = 73, 58.40%, **Supplementary Fig. 6 and Supplementary Table 13**), including ammonia-oxidizing bacteria (*Nitrosomonadaceae*), methylotrophs (*Methyloligellaceae*), and sulfate-reducing bacteria (*Desulfobacterales*). These results suggest that their extensive reliance on multiple quorum sensing systems allows these hub microorganisms to facilitate broad interspecies communication and interactions within the cold seep ecosystem.

### Quorum sensing regulates functionally key microbial groups in cold seeps

Significant correlations between quorum sensing genes and key metabolic genes suggest that quorum sensing impacts key biogeochemical functions such as anaerobic methane oxidation, sulfate reduction, nitrogen fixation, and reductive dehalogenation (*P* < 0.05, **Fig. 4a-d and Supplementary Table 14**). These processes, mediated by genes like *mcrA*, *dsrA*, *nifH*, and *rdhA*, are vital for energy production and resource acquisition in cold seep microbial communities^50–53^. Positive correlations between *dsrA* and AHL synthase gene *hdtS*, AHL-lactonase gene, and receptor gene *luxN* (**Fig. 4a**) suggest that quorum sensing may influence sulfate reduction, as previously observed in *Desulfovibrio vulgaris* and *Desulfobacterium corrodens* in marine environments^54^. Similarly, a significantly positive correlation between *mcrA* and the PQS synthase gene *pqsE* (**Fig. 4b**) hints that PQS signaling may be involved in regulating methane oxidation. Additionally, *rdhA* showed positive correlations with multiple quorum sensing genes, including AI-2 transporter genes, AHL degrading enzyme genes, and DSF synthase gene *rpfF* (**Fig. 4c**), indicating that quorum sensing might influence the metabolism of organohalides. Notably, some studies have demonstrated that organohalides can interfere with quorum sensing signals, which may serve as a defense mechanism against microbial colonization on algal surfaces^53,55^. Furthermore, *nifH* was positively correlated with *pqsE* but negatively correlated with multiple quorum sensing genes (**Fig. 4d**), suggesting that nitrogen-fixing microorganisms may engage less frequently in quorum sensing-mediated communication.

**Figure 4.**
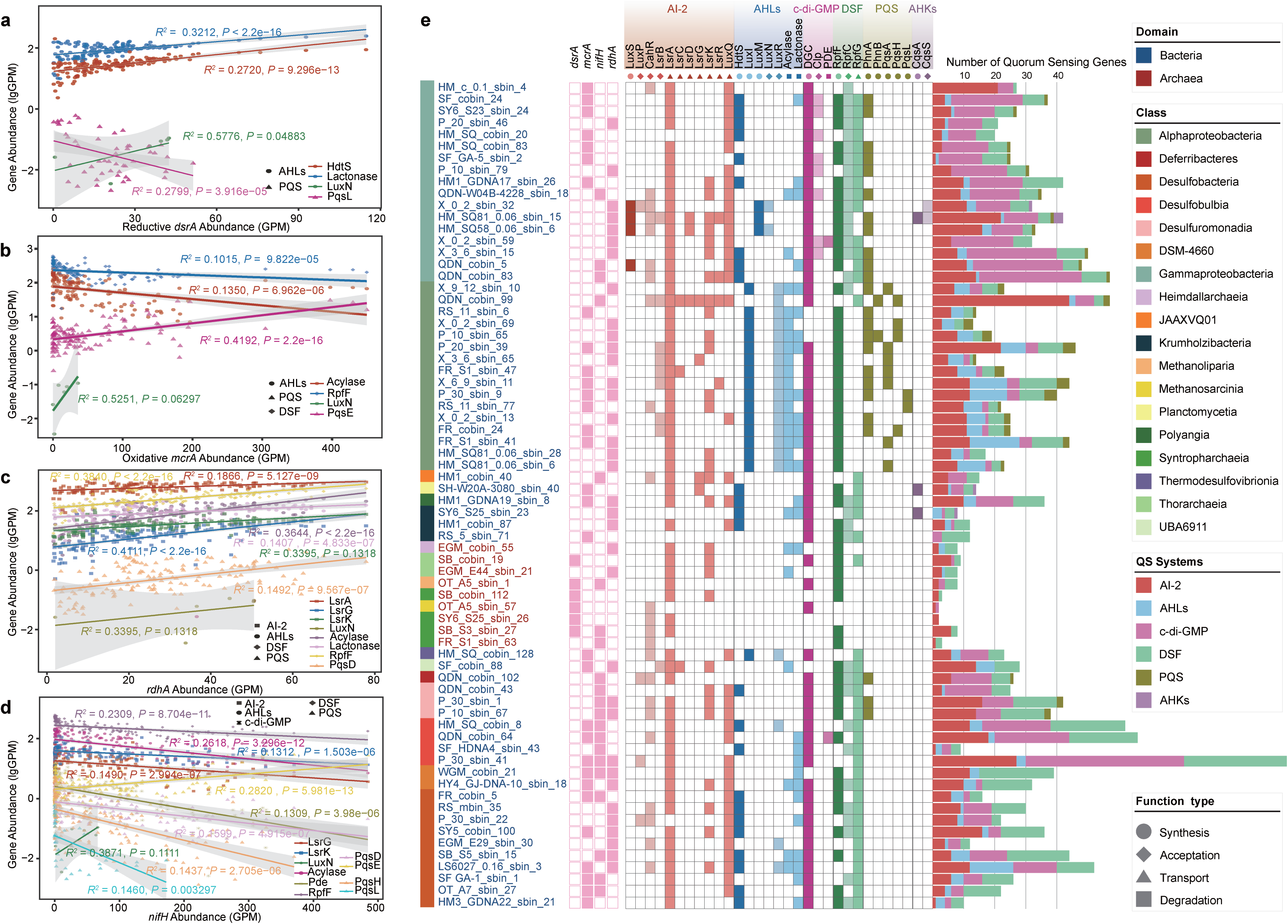
Relationship between quorum sensing genes and key metabolic genes in cold seeps and distribution of quorum sensing genes among functionally key microbial groups. **(a–d)** Correlation between quorum sensing gene abundance and key metabolic genes in cold seep sediments, including (a) *dsrA* (sulfate reduction-related), (b) *mcrA* (methane oxidation-related), (c) *rdhA* (dehalogenation-related) and (d) *nifH* (nitrogen fixation-related). Linear regression lines, *P*-values and adjusted *R²* (Radj²) values are provided, indicating the strength and significance of the relationships. **(e)** Presence of key metabolic genes and various quorum sensing systems in functionally key microbial groups (ANME, SRB, diazotrophs and organohalide reducers). Different phyla are denoted by the colored bars in the first column; the presence of quorum sensing genes in functionally key microbial groups is indicated by colored squares, with distinct colors representing various types of quorum sensing systems. Stacked bar charts in the last column represent the proportion of each quorum sensing system within these microorganisms. Detailed data are provided in **Supplementary Tables 14-15**.

A detailed analysis of quorum sensing genes distribution across 1,084 functionally key microbial groups in cold seeps, including ANME, SRB, diazotrophs, and organohalide reducers, provides further insights into the regulatory role of quorum sensing in cold seep microbial processes. In ANME, including ANME-1 and ANME-3, quorum sensing genes were predominantly associated with AI-2 and DSF systems, focusing on the synthesis and transport of signaling molecules (**Fig. 4e and Supplementary Table 15**). In contrast, SRB exhibited a broader distribution of quorum sensing genes across all systems, with particularly high levels of c-di-GMP-related genes, AHL synthase genes, and DSF-related genes (**Fig. 4e and Supplementary Table 15**). Notably, 19.70% of identified *clp* genes, which encode global transcription factor receptors^56^, were found in SRB. This implies that the presence of multiple quorum sensing signals enhances the communicative flexibility of these microorganisms, potentially facilitating extensive intercellular communication in cold seep ecosystems^48^.

Diazotrophs (nitrogen-fixing microorganisms) also encoded various quorum sensing genes, particularly DSF-related ones (**Fig. 4e and Supplementary Table 15**). Interestingly, five of these MAGs contained genes for all six quorum sensing systems, suggesting that these microorganisms engage in extensive communication and coordination within the community, likely enhancing resource utilization and metabolic efficiency during nitrogen fixation. Organohalide reducers exhibited unique quorum sensing profiles, with a high prevalence of genes related to AI-2 systems, particularly transporters, and a greater abundance of AHK-related genes compared to other functional groups (**Fig. 4e and Supplementary Table 15**). Based on their quorum sensing preferences, organohalide reducers were grouped into three categories (**Supplementary Fig. 7**): (i) those that primarily used c-di-GMP with a high presence of DGC genes and *clp*, (ii) those focusing on AHLs, and (iii) those predominantly employing DSF signaling. Overall, the findings suggest that microorganisms involved in key metabolic pathways are likely regulated by a synergistic action of multiple quorum sensing systems.

### Cold seep microbial behaviors and biogeochemical processes are modulated through c-di-GMP PDE and three key receptor types

To explore the microbial behaviors and biogeochemical processes potentially regulated by key quorum sensing-related proteins in cold seep ecosystems, we examined the genetic characteristics of key quorum sensing genes, focusing on the c-di-GMP-degrading enzyme PDE and three receptor types: CahRs, RpfC/G two-component systems, and LuxR. PDE degrades c-di-GMP, thereby increasing the proportion of free c-di-GMP receptor Clp^57^. Free Clp can bind to various DNA fragments, enabling microbes to use a “mobile-attack” strategy during bacterial competition^57^. Interestingly, some PDEs contain both synthesis (GGDEF) and degradation (EAL) domains^58,59^ (**Fig. 5a**), suggesting that certain microorganisms can regulate both the synthesis and degradation of c-di-GMP simultaneously. This trait allows for dynamic control of critical processes such as host-microbe symbiosis, biofilm formation, motility, and virulence, all through the modulation of c-di-GMP levels via the regulation of PDE expression^60,61^.

**Figure 5.**
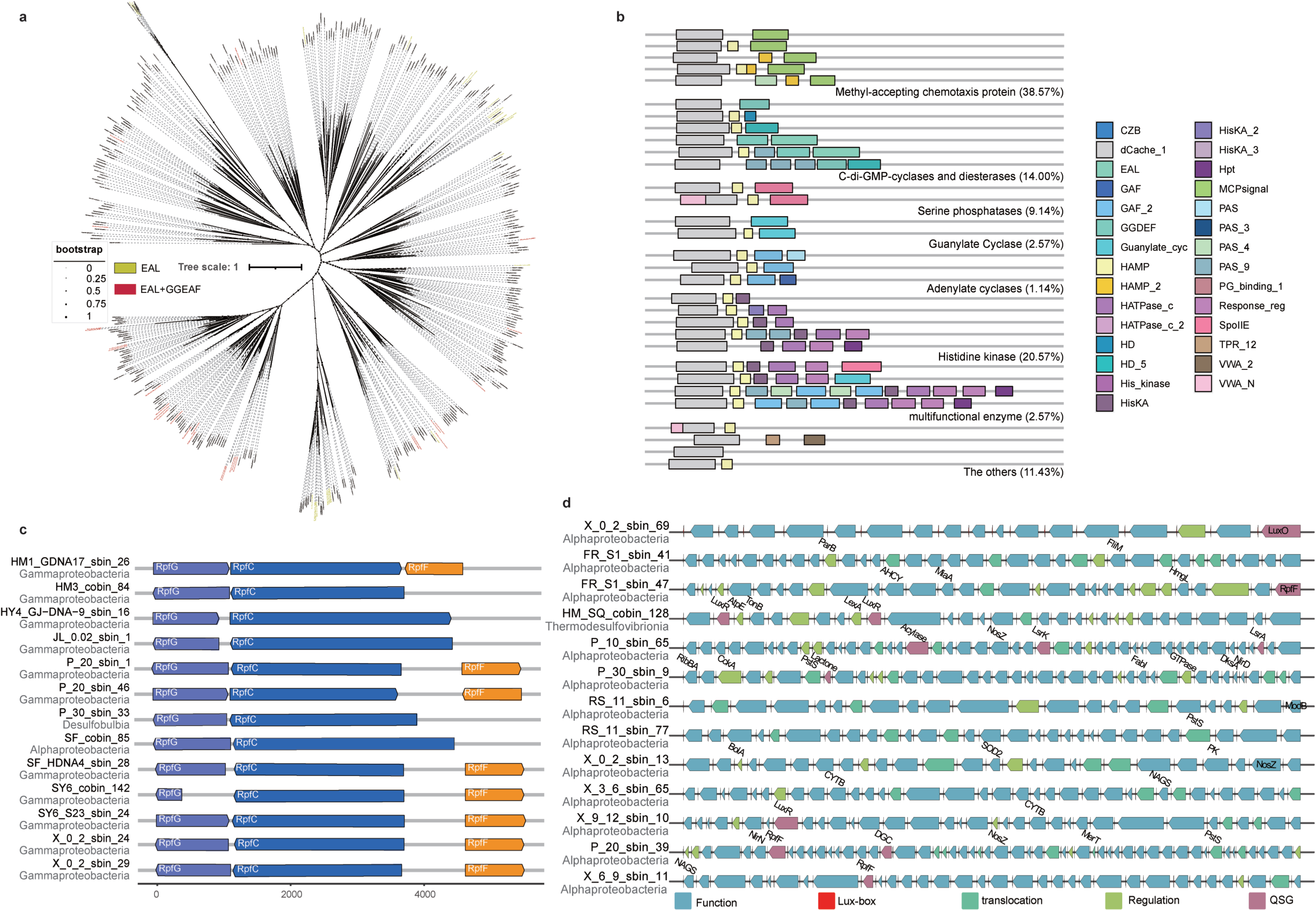
Structural characteristics and functional analysis of quorum sensing genes in cold deep microorganisms. **(a)** Maximum-likelihood phylogenetic tree of phosphodiesterases (PDEs) possessing EAL motifs and HD-GYP conserved sites derived from cold seep microorganisms. PDEs are grouped based on functional domain characteristics, indicated by different label colors. The scale bar represents the average number of substitutions per site. **(b)** Schematic representation of the domain structures of 350 CahR-type receptor proteins. Detailed information on CahR-type receptor protein domain predictions is provided in **Supplementary Table 16**. **(c)** Illustration of 13 genomic fragments containing the RpfC-G two-component system within the DSF signaling pathway. Genes are depicted as arrows, with different colors representing various functions (e.g., sensor kinase RpfC, response regulator RpfG). **(d)** Visualization of predicted *lux* box sequences and their potentially regulated downstream genes in 13 MAGs containing LuxI-R pairs. Genes are colored based on their functions: functional genes (blue), transport-related genes (dark green), regulatory genes (light green), and quorum sensing genes (purple). Detailed information is provided in **Supplementary Table 17.**

We identified 350 CahR-type receptors across cold seep microorganisms, with 318 of these receptors associated with signal transduction proteins that may function as extracellular signal sensors (**Supplementary Table 16**). The CahR receptors, containing the dCACHE_1 domain, serve as sensors of extracellular signals from transmembrane proteins like methyl-accepting chemotaxis proteins (MCPs), histidine kinases (HKs), c-di-GMP synthases, and phosphodiesterases (GCDs), mediating both intraspecies and interspecies communication^39^. Among these, MCPs (38.57%) and HKs (20.57%) were the most prevalent signal transduction proteins regulated by AI-2 (**Fig. 5b**). MCPs and HKs regulate microbial motility and chemotaxis, enabling microorganisms to relocate towards more favorable environments^62^. Additionally, HKs mediate the metabolism of citrate and malate by initiating phosphorylation cascades^63–65^. A total of 150 CahRs were identified among microorganisms involved in key metabolic processes, with *Pseudomonadota* (n = 50) and *Desulfobacterota* (n = 40) containing the highest numbers of CahRs.

We also identified 13 RpfC/G two-component systems involved in DSF signaling (**Fig. 5c**). The RpfC/G two-component system is essential for DSF signaling, where DSF is synthesized by RpfF, detected by the sensor kinase RpfC, and transduced through the response regulator RpfG^66,67^. This system may be used by microorganisms to regulate the synthesis of extracellular enzymes and polysaccharide virulence factors^66^. Additionally, RpfG can degrade c-di-GMP, similar to PDE, increasing the ratio of c-di-GMP-free Clp, thereby driving the expression of hundreds of genes^68,69^. This highlights the collaborative and interconnected nature of quorum sensing systems in cold seep microorganisms.

A total of 15 LuxI-R pairs were identified from 13 MAGs involved in functionally key microbial groups, with 437 *lux* box-like 20 bp palindromic sequences associated with these LuxI-R pairs (**Fig. 5d and Supplementary Table 17**). The *lux* box is the most specific binding site of the LuxR/AHL complex^70^. Analysis of these *lux* box sequences suggests that LuxR is involved in the regulation of genes associated with elemental cycles, including *nosZ* (encoding nitrous oxide reductase)^71^, *nirN* (encoding nitrate reductase)^72,73^, *norD* (encoding nitric oxide reductase)^73^, and *soxY* (encoding sulfur-oxidizing protein)^74^. LuxR also regulates other quorum sensing genes, such as AHL-lactonase gene, *rpfF*, and DGC gene, highlighting the complex interactions between different quorum sensing systems. Additionally, LuxR potentially influences the molybdenum (Mo) cycle, which is enriched in cold seep sediments^75^, by regulating Mo transport gene *modB*^76,77^. The LuxR may also influence cold seep microorganisms by affecting microbial physiology. For instance, LuxR could indirectly control microbial motility by regulating the expression of *fliM*, a gene involved in flagellum production and assembly^78^. Furthermore, LuxR might reduce virulence and inhibit biofilm formation via *fabI*^79^. These findings underscore the central role of LuxR in facilitating cold seep microbial survival and ecological interactions.

### Heterologous expression and sediment metabolomics confirmed occurrence of quorum sensing in cold seeps

Our results provide strong evidence for the presence of quorum sensing systems in cold seep environments. To assess the functional activity of *luxI* genes, we performed heterologous expression experiments with five *luxI* genes containing conserved active sites (R25, F29, W35, D46, R70, E101, R104; **Fig. 6a**), to evaluate their ability to synthesize AHLs. Recombinant *E. coli* strains expressing four of these genes produced blue pigments on indicator plates, while no pigment was observed in controls containing only the pET-28a vector (**Fig. 6b–6c**). This indicates that these four *luxI* genes are functional and capable of synthesizing AHLs in a heterologous system, confirming their role in quorum sensing. These active *luxI* genes belong to representatives from candidate genera *JAADFS01*, *UBA3077*, and *JAACFB01* within *Nitrospirota* and *Pseudomonadota* (**Supplementary Table 8**). Further analysis of P_20_sbin.39*-luxI* revealed its co-localization with P_20_sbin.39*-luxR*, forming a LuxI-LuxR pair in P_20_sbin.39. Molecular docking analysis demonstrated that the P_20_sbin.39-LuxR complex exhibited binding affinities of −5.15 kcal/mol, −5.13 kcal/mol, −5.28 kcal/mol, and −6.41 kcal/mol with C_10_H_15_NO_4_, C_12_H_21_NO_3_, C_14_H_25_NO_3_, and C_16_H_27_NO_4_, respectively (**Fig. 6d**). These molecules are likely autoinducers, which trigger gene expression through a self-talk mechanism typical of LuxI-LuxR quorum sensing systems^48^.

**Figure 6.**
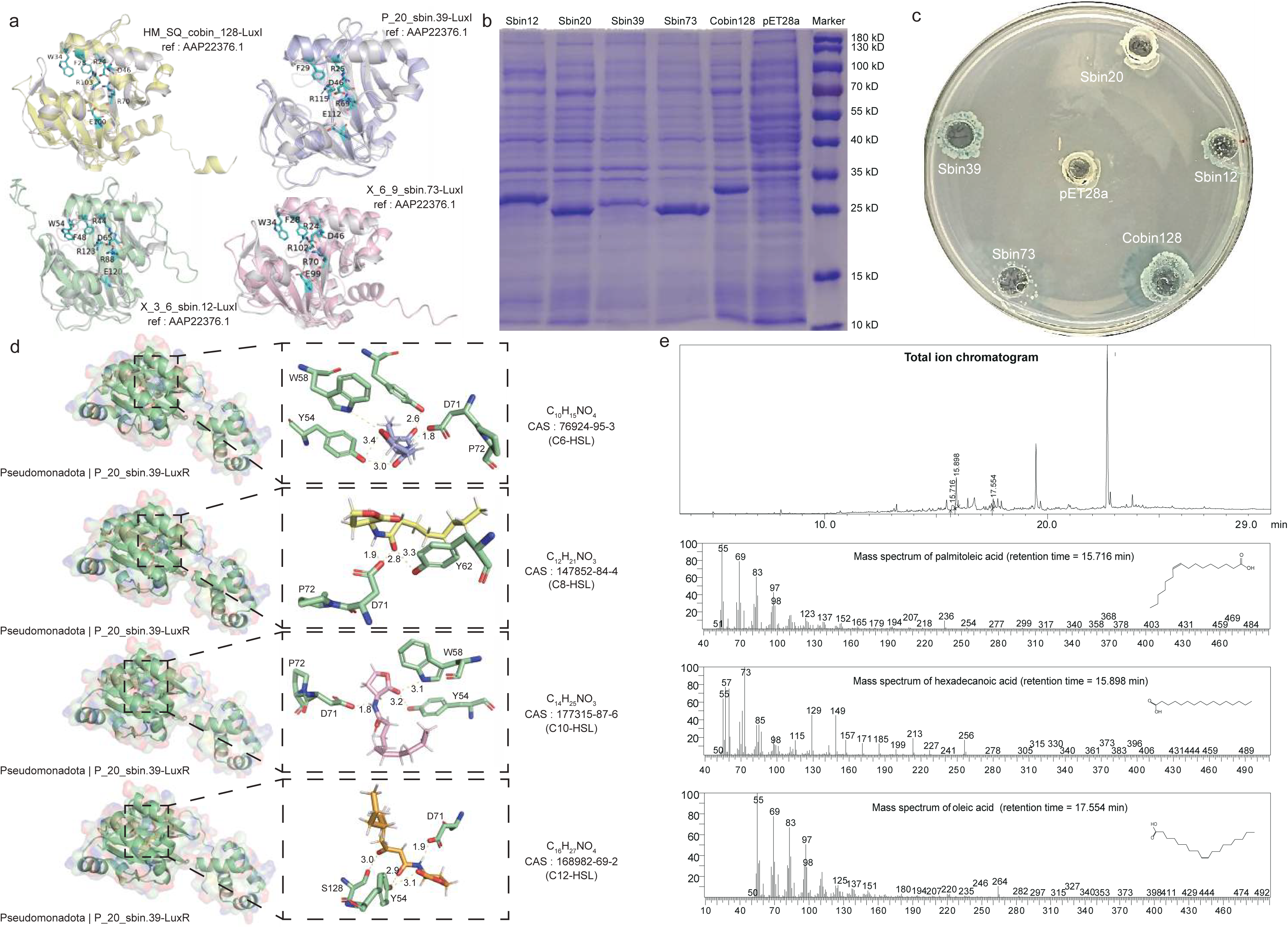
LuxI expression and bioassay, and detection of quorum sensing-related compounds in cold seep sediments by GC-MS. **(a)** Overlay of the active sites of cold seep-derived active LuxI (colored) onto known LuxI structures (gray). The active site is shown sticks. **(b)** Expression of LuxI. SDS-PAGE analysis of LuxI in recombinant *E. coli* BL21, with molecular sizes (kDa) indicated. **(c)** Bioassay of LuxI with the reporter strain *A. tumefaciens* A136 for AHL activity of LuxI. Extracts from recombinant *E. coli* BL21 strains carrying X_3_6_sbin.12-*luxI*, P_20_sbin.39-*luxI*, X_6_9_sbin.73-*luxI*, and HM_SQ_cobin_128-*luxI* developed blue pigments, whereas the control strain, *E. coli* BL21 with pET28a, showed no blue pigments. **(d)** The protein structures of LuxR and their molecular docking with AHLs. Overall (left) and local magnification (right) of molecular docking of P_20_sbin.39-LuxR with AHLs. The residues forming polar contacts (indicated by yellow dashed line) of P_20_sbin.39-LuxR are shown in green sticks. The ligands C_10_H_15_NO_4_, C_12_H_21_NO_3_, C_14_H_25_NO_3_, and C_16_H_27_NO_4_ are represented by blue, yellow, pink, and orange sticks. **(e)** GC-MS analysis of quorum sensing-related compounds in cold seep sediments. The upper part shows the total ion chromatogram, and the corresponding peaks for hexadecanoic acid, palmitoleic acid, and oleic acid in cold seep sediment are shown below the chromatogram.

To further confirm the occurrence of quorum sensing in cold seep environments, we used gas chromatography-mass spectrometry (GC-MS) to detect autoinducers in cold seep sediments. Quorum sensing typically operates at very low concentrations (as low as 1 pM)^80^, which makes in situ detection of autoinducers challenging. Although we did not detect autoinducers directly, we successfully identified several quorum sensing-related compounds, including hexadecanoic acid, oleic acid, and palmitoleic acid (**Fig. 6c and Supplementary Fig. 8**). Hexadecanoic acid has been shown to inhibit CviR-mediated quorum sensing^81^, while oleic acid blocks the binding of AHLs by interacting with AbaI and suppresses the AI-2-signaling pathway^82,83^. Furthermore, palmitoleic acid interferes with the DSF-mediated signaling pathway, inhibiting quorum sensing^84^. GC-MS-based metabolomic analysis provides indirect evidence that cold seep sediments contain quorum-sensing compounds, which may modulate microbial communication pathways.

## Conclusions

In summary, our study provides compelling evidence of the widespread presence and functional significance of quorum sensing systems in cold seep environments. We identified six major quorum sensing systems, including AI-2, AHLs, DSF, c-di-GMP, PQS, and AHKs, across diverse microbial communities in cold seep sediments. These systems displayed depth-dependent patterns and exhibited considerable variability between different cold seep types, suggesting that microorganisms in these ecosystems have evolved flexible and highly adaptable communication strategies to respond to their unique environmental conditions. The integration of multiple quorum sensing pathways allows microorganisms to coordinate both intraspecies and interspecies interactions, enhancing their ability to thrive in these challenging environments. Our findings highlight the central role of quorum sensing in regulating key microbial behaviors, including motility, biofilm formation, and stress responses, which are essential for survival in the nutrient-limited and chemically dynamic cold seep habitats. Furthermore, quorum sensing appears to influence critical biogeochemical cycles, such as carbon, nitrogen, and sulfur transformations, reinforcing the importance of microbial communication in sustaining ecosystem processes. Overall, our research underscores the complexity of microbial communication in the deep biosphere, revealing how multiple quorum sensing systems enable microorganisms to thrive and contribute to the ecological stability of cold seep ecosystems. These insights into microbial adaptation and signaling may have broader implications for understanding microbial interactions in other extreme environments and could pave the way for future studies on the biogeochemical roles of microorganisms in the deep ocean.

## Materials and Methods

### Metagenomic and metatranscriptomic data processing

A total of 173 metagenomes, 33 metatranscriptomes and 18 metabolomes were obtained from 17 geographically diverse cold seep sites^32,85^ (**Supplementary Fig. 1**). Detailed information about sampling sites, sediment depths and sequencing protocols is available in our previous publications^32^. Briefly, to construct a non-redundant gene catalog, metagenomic sequencing data was quality controlled to produce clean reads. Assembly was then performed using MEGAHIT (v1.1.3; parameters: --k-min 27 --kmin-1pass --presets meta-large)^86^. Protein-coding sequences were predicted for these assembled contigs using Prodigal (v2.6.3; parameter: -meta)^87^ and sequences were clustered at 95% amino acid identity (AAI)^88^, resulting in a non-redundant gene catalog with 147,289,169 genes. Relative abundance of these genes was quantified using Salmon (v1.10.2)^89^ and normalized to GPM. For constructing the non-redundant MAG catalog, assembled contigs exceeding 1 kb in length were subjected to subsequent binning, followed by dereplication of MAGs using dRep (v3.4.0; parameters: -comp 50 -con 10)^90^ for clustering at a 95% average nucleotide identity (ANI) threshold, ultimately yielding a total of 3,813 MAGs. Taxonomic classification of MAGs was performed using GTDB-Tk (v2.1.1) with reference to the GTDB R207 database^91^. Phylogenomic trees for archaea and bacteria were constructed using the maximum-likelihood method implemented in FastTree (v2.1.11, default parameters)^92^ and visualized with iTOL (v6)^93^. A sankey plot was generated using Pavian (https://fbreitwieser.shinyapps.io/pavian/)^94^ to visualize data flow between taxonomic groups. Functional annotation of MAGs was performed using DRAM (v1.3.5; parameter: --min_contig_size 1000), utilizing KEGG, Pfam, MEROPS and dbCAN databases^95^.

For metatranscriptomic data, quality filtering was conducted using the Read_QC module within the metaWRAP pipeline (v1.3.2, parameters: --skip-bmtagger)^96^. Ribosomal RNA (rRNA) was removed from the quality-filtered reads using SortMeRNA (v2.1, default parameters)^97^. Transcript abundance of non-redundant genes were quantified using Salmon (v1.9.0, parameters: -validateMappings -meta)^89^, with results normalized to TPM.

### Identification of quorum sensing genes in MAGs and non-redundant gene catalog

To identify quorum sensing genes in MAGs and the non-redundant gene catalog, we used the QSP database and searched for potential quorum sensing sequences using HMMER (v3.2.1, parameters: cut_ga)^31^.

For MAGs, we focused on sequences related to key quorum sensing systems, including AI-2, AHLs, DSF, c-di-GMP, PQS, and AHKs. Phylogenetic analysis based on reference sequences and alignment results from HMMER searches were used to identify these quorum sensing genes. Sequences were aligned using MAFFT (v7.505, default settings)^98^ and trimmed using TrimAL (v1.4.1, default settings)^99^. Maximum-likelihood phylogenetic trees were constructed using FastTree (v2.1.11, default settings)^92^ and visualized using iTOL (v6)^93^. Conserved active site alignment was performed for identified HdtS, Acylase, Lactonase, DGC, PDE, Clp, RpfF, RpfG, PqsA and PqsD, followed by visualization of the results using MAFFT (v7.505, default settings)^98^, Jalview^100^ and WebLogo3 (https://weblogo.threeplusone.com/). Protein domains of LsrA, LuxQ, and CahR were annotated using the Pfam database (http://pfam.xfam.org/), while domains of Clp and RpfC were predicted using the SMART database^101^. Deep learning approach AlphaFold2 was used to predict three-dimensional structures of LuxI, LuxS, LuxP and PqsL^102^. Structural similarity clustering results, obtained from Foldseek (https://cluster.foldseek.com/)^103^, were assessed using Template Modeling Score (TM-score) and Root Mean Square Deviation (RMSD), with sequences considered valid if they met criteria of TM-score > 0.5 and RMSD < 1^104^. The comparative results were visualized using PyMOL^105^.

For the non-redundant gene catalog, we focused on a large number of sequences and validated quorum sensing genes such as CahR, LsrA, LsrK, LuxR, DGC, PDE, RpfF, RpfG, PqsD, and PqsL using the same methods outlined above. Other genes were validated through phylogenetic tree construction with reference sequences (software and parameters as described above). Detailed methods for the screening and validation of quorum sensing genes are provided in **Supporting Methods**, **Supplementary Figs. 9-19 and Supplementary Table 1**.

### Identification of viral genomes containing quorum sensing genes

The prediction and functional annotation of viral sequences followed methods from our previous research^85^. In brief, potential viral sequences were identified from contigs (length > 10 kb) assembled from 173 cold seep samples using geNomad (v1.7.0, parameters: end-to-end)^106^. Sequences with a completeness of 50% or higher were clustered into species-level viral operational taxonomic units (vOTUs) using CheckV (95% nucleotide identity and at least 85% aligned fraction)^107^, resulting in 17,051 vOTUs. Taxonomic assignments of vOTUs were performed using geNomad (v1.7.0). Open reading frames (ORFs) within each vOTU were predicted using Prodigal (v2.6.3, parameter: -p meta)^87^. Functional annotation of viral sequences was conducted using the virus model in DRAM (v1.4.0)^95^. Potential quorum sensing genes in viral genomes were identified using HMMs from the QSP database, following aforementioned manual screening methods. The downstream and upstream functional genes associated with viral quorum sensing genes were visualized using Chiplot (https://www.chiplot.online/).

### Correlation network based on the relative abundance of quorum sensing genes in cold seep microorganisms

To explore potential interaction patterns between quorum sensing genes and cold seep microbial genomes, a correlation network based on the relative abundance of quorum sensing genes, identified from the non-redundant gene catalog, and MAGs was constructed. Only MAGs detected in more than 20% of samples were included to minimize false-positive correlations. Spearman’s rank correlation coefficients were calculated in R (v4.3.1) based on the relative abundance of quorum sensing genes and MAGs. Only significant correlations (|*R*| ≥ 0.5, *P* < 0.05) were retained. Correlation networks were visualized using Cytoscape (v3.10.1)^108^ and Gephi (v0.10.1)^109^, where nodes represent quorum sensing genes and MAGs, and edges reflect the strength and significance of the correlations.

### Identification of *mcrA*, *dsrA*, *nifH* and *rdhA*

For identification of *mcrA*, *dsrA,* and *nifH,* previous studies were referenced^85^. Briefly, *mcrA* and *dsrA* were identified using METABOLIC (v4.0)^110^ and DRAM (v1.4.0)^95^, and then *dsrA* was further detected using DiSco (v1.0.0)^111^. The *nifH* was analyzed using DIAMOND blastp (v2.0.8, parameters: --id 65)^112^ against NCycDB^113^ and the Greening Lab database (https://bridges.monash.edu/collections/_/5230745). The candidate *nifH* sequences were then extracted using HMMER (v3.3.2) with the TIGRFAM model TIGR01287.1. For identification of the dehalogenation gene *rdhA*, the methodologies referenced in prior studies were employed^80^. The *rdhA* was analyzed using DIAMOND blastp (v2.0.15.153)^112^ against the Reductive Dehalogenase Database (https://rdasedb.biozone.utoronto.ca/)^114^. The Hmmsearch tool in HMMER (v3.3.2, E-value < 1×10^−10^) was applied using HMM profile “PF13486” and NCBI HMM accession “TIGR02486”. Protein length (> 300 aa) and presence of two conserved iron-sulfur (Fe-S) motifs (CXXCXXCXXXCP, CXXCXXXCP) were manually verified.

### Prediction of AHL-regulated genes in cold seep microorganisms

Thirty LuxI-R pairs were identified through gene co-localization analysis. For functionally key microbial groups containing LuxI-R pair, the upstream region (300 bp) of each ORF was extracted. Putative *lux* box sequences in these upstream regions were predicted using MEME-FIMO^115^, trained with a library of 54 highly conserved and previously reported *lux* box sequences. Three criteria were applied during the prediction process^116^: (i) 20 bp nucleotide length, (ii) imperfect palindromic sequence, and (iii) a default *P*-value of < 0.0001. DRAM (v1.3.5) was used to functionally annotate downstream genes of the predicted *lux* box sequences.

### Construction of three-dimensional Structures and molecular docking

AlphaFold2 was used to predict the three-dimensional structure of the P_20_sbin.39-LuxR sequence^102^. To evaluate its binding potential, we performed molecular docking using LeDock (v1.0; https://www.lephar.com) with a root-mean-square deviation (RMSD) cutoff of 1.0 and 20 binding conformations. Four different acyl-homoserine lactones (HSLs), each with varying carbon chain lengths, were selected as ligands. The Autoinducer binding domain of P_20_sbin.39-LuxR was predicted using PyMOL^105^ based on the experimentally validated structure of LuxR (PDB: 3SZT, from *Pseudomonas aeruginosa*). A binding box was generated using the GetBox PyMOL plugin (extension = 5.0; https://github.com/MengwuXiao/GetBox-PyMOL-Plugin), and the docking results were visualized.

### LuxI protein expression and bioassay

X_3_6_sbin.12-*luxI*, P_20_sbin.39-*luxI*, X_6_9_sbin.73-*luxI*, HM_SQ_cobin_128-*luxI* and XST_SQ83_0.10_sbin_20_*luxI* synthesis and plasmid construction were performed by Sangon Biotech. The amplified genes were inserted into the pET-28a expression vector and transformed into *Escherichia coli* BL21 cells. A 1% inoculum was introduced into fresh Luria-Bertani (LB) medium containing kanamycin (50 µg/ mL) and introduced at 20°C with shaking at 150 rpm. When the culture reached an optical density at 600 nm (OD_600_) of 0.4-0.6, isopropyl-β-D-thiogalactopyranoside (IPTG) was added to a final concentration of 0.2 mM, and the culture was incubated overnight. Cells were harvested and stored for further use. The harvested cells were resuspended in 50 mM phosphate buffer and lysed by ultrasonication. The lysate was analyzed by 10% SDS-PAGE to verify the expression of the target gene.

To assess the production of AHLs by the LuxI-expressing strains, *Agrobacterium tumefaciens* A136 was used as biosensors^117^, which produces a blue pigment in thepresence of AHLs. The A136 strain was cultured overnight at 30 °C, and 100 μL of the fresh culture was evenly spread onto LB agar plates using the pour plate method. After incubation at 30 °C for 24 hours, a layer of water agar containing a final concentration of 40 µg/L of X-gal was added on top. Once the water agar solidified, wells were created, and 50 μL of the expressing culture was added to each well, which were then sealed with water agar. After 24 hours of incubation at room temperature, the activity was assessed.

### Analysis of metabolites using GC-MS

Metabolite extraction and analysis were performed following a modified protocol based on previous studies^118^. Briefly, 10 g of cold seep sediments were placed in 40 mL of ethyl acetate and extracted by shaking at 120 rpm for 2 h, with centrifuge tubes gently swirled every 15 min. After extraction, centrifuge tubes were centrifuged at 2,000 rpm for 20 min, and the supernatant was collected. The sediment underwent a second extraction with fresh 40 mL of ethyl acetate under the same conditions. Supernatants from both extractions were combined and concentrated to 1 mL using a rotary evaporator at 50 °C. Purification was carried out using a three-layer solid-phase extraction (SPE) column, composed of anhydrous sodium sulfate (1.0 g), silica gel (0.5 g) and anhydrous sodium sulfate (0.5 g). The SPE column was pre-washed with 5 mL of *n*-hexane before sample application. The concentrated extract was added to the column and firstly eluted with 8 mL *n*-hexane, and then eluted with 8 mL of ethyl acetate. The ethyl acetate eluate was collected, concentrated to near dryness using a rotary evaporator at 50 °C, and reconstituted to 1 mL with ethyl acetate. The final sample was filtered through a 0.22 µm organic solvent-resistant membrane filter for GC-MS analysis. GC-MS analysis was conducted using an Agilent HP-5ms Capillary GC column (30 m × 0.25 mm i.d. × 0.25 µm film thickness) on a QP2010 Ultra instrument (Shimadzu). The injection port temperature was set to 290 °C. The oven temperature program started at 100 °C, held for 1 min, ramped to 150 °C at 35 °C/min, then to 280 °C at 25 °C/min, where it was held for 6 min, and finally increased to 290 °C, holding for 2 min. The mass spectrometer operated in selected ion monitoring (SIM) mode with an electron ionization energy of 70 eV. The ion source and quadrupole temperatures were set at 230 °C and 150 °C, respectively. Hexadecanoic acid, palmitoleic acid, and oleic acid were identified by comparing the measured compounds with mass spectral libraries and the spectra of authentic standards.

### Statistical analyses

Statistical analyses were conducted using R (v4.2.3). The Kolmogorov-Smirnov test was applied to assess normality, while Levene’s test was used to evaluate homogeneity of variance. To compare the correlations between relative abundances of different signaling systems and quorum sensing genes, the Kruskal-Wallis rank-sum test was applied. Differences in quorum sensing systems across depths and cold seep types were evaluated using the Wilcoxon test. Pearson correlation and linear regression analyses were conducted to evaluate the relationships between quorum sensing genes and key metabolic genes, including *mcrA*, *dsrA*, *nifH,* and *rdhA*.

### Code and data availability

The non-redundant gene and MAG catalogs from 173 cold seep metagenomes are available in Figshare (https://doi.org/10.6084/m9.figshare.22568107). The phylogenetic tree based on amino acid sequences of quorum sensing proteins, along with corresponding protein structure files, can be accessed at Figshare (https://doi.org/10.6084/m9.figshare.27268377). This study did not generate custom code, and the mentioned tools for data analysis were applied using default parameters unless specified otherwise.

## Supporting information

QS-inf

## Acknowledgements

This work was supported by National Natural Science Foundation of China (No. 92351304, No. 42376115 and No. 32170121), Natural Science Foundation of Fujian Province (No. 2023J06042), Natural Science Foundation Project of Xiamen City (No. 3502Z202373076), and Scientific Research Foundation of the Third Institute of Oceanography, Ministry of Natural Resources (No. 2022025 and No. 2023022). M. R.-B. is funded by the Israeli Science Foundation (ISF) grant 1359/23. We thank Bin Wei and Gangao Hu for their assistance with metabolomic analyses. We are also grateful to Lingwei Ruan, Hong Shi, and Zengtian Gu for their contributions to heterologous expression.

## Author contributions

X.D. conceived and designed the study. J.P. and X.L. performed the omics analyses. Q.M., J.W., N.M., and J.P. conducted the protein expression and enzymatic activity assays. R.C. contributed to the metabolomic data analysis. Y.P., Y.H., J.L., C.L., and M. R.-B. actively participated in discussions and data interpretation. J.P., X.L., and X.D. wrote the manuscript with input from all authors.

## Competing interests

The authors declare no competing interests.

## Notes

### Competing Interest Statement

The authors have declared no competing interest.

