## Supplementary material for "Diverse quorum sensing systems regulate microbial communication and biogeochemical processes in deep-sea cold seeps": QS-inf

Supporting Methods

The identification of quorum sensing genes from the non-redundant gene catalog with 147 million genes and 3,813 species-level representative metagenome-assembled genomes (MAGs) is based on the quorum sensing-related proteins (QSP) database^1^, combined with multiple screening approaches. Detailed information is provided below (**Supplementary Table 1**):

**Identification of quorum sensing genes associated with the AI-2 system**

For the 3,813 species-level representative MAGs, phylogenetic trees of the amino acid sequences of LsrC, LsrD, LsrG, LsrR, LuxS, CahR, LuxP, and LsrB were constructed to validate their phylogenetic clades against reference sequences **(Supplementary Fig. 9)**. Sequences were aligned using MAFFT (v7.505, default settings)^2^ and trimmed using TrimAL (v1.4.1, default settings)^3^. Maximum-likelihood trees were constructed using FastTree (v2.1.11, default settings)^4^, and all trees were visualized and beautified using iTOL (v6)^5^. Candidate LuxP and LuxS sequences that clustered poorly with reference sequences further validated using AlphaFold2^6^. These structures were compared with the structural similarity clustering results obtained from Foldseek (https://cluster.foldseek.com/)^7^, and assessments were performed using the Template Modeling Score (TM-score)^8^ and Root Mean Square Deviation (RMSD)^9^. Sequences were considered valid if they met the criteria of TM-score > 0.5^10^ and RMSD < 1 **(Supplementary Fig. 9i-9j)**. The comparative results were visualized using PyMOL^11^. Additionally, LsrR sequences were extracted using HMMER (v3.3.2) with the profile “PF04198”^12^ and LsrK sequences using profiles “PF00370” and “PF02782”^13^. For CahR, sequences were extracted using HMMER (v3.3.2) with the profile “PF02743”^14^. Candidate CahR and LsrA sequences were further analyzed for domain structure using the Pfam database (https://www.ebi.ac.uk/interpro/), confirming the ATP-binding cassette domain in LsrA^15^.

For the non-redundant gene catalog with 147 million genes, the CahR, LsrA and LsrK were screened using the same method as the genome-based analysis. LuxS, LuxP, LsrB, LsrC, LsrD, LsrG, LsrR and LuxQ were identified based on their phylogenetic clades with reference genes **(Supplementary Fig. 10)**. The software and parameters used for constructing the phylogenetic tree are the same as those used in the genome-based analysis.

**Identification of quorum sensing genes associated with the AHLs system**

For the 3,813 species-level representative MAGs, phylogenetic trees of HdtS, LuxM, LuxI, LuxN, AHL-acylase and AHL-lactonase were constructed to validate their phylogenetic clades against reference sequences **(Supplementary Fig. 11)**. Sequences were aligned using MAFFT (v7.505, default settings)^2^ and trimmed using TrimAL (v1.4.1, default settings)^3^. Maximum-likelihood trees were constructed using FastTree (v2.1.11, default settings)^4^, and all trees were visualized and beautified using iTOL (v6)^5^. For HdtS, sequences containing conserved motifs NHQS and PEGTR were retained ^16^. For LuxR, sequences containing both the autoinducer-binding domain (PF03472) and the GerE domain (PF00196) were extracted^17^. Sequences of LuxI were identified using HMMER (v3.3.2) with the profile “PF00765”^17^. For AHL-acylase, sequences containing the key amino acid Ser217 were retained^18^, and sequences containing the conserved motif HXHXDH-60aa-H were selected as AHL-lactonase^19^. Conserved sites were visualized using MAFFT (v7.505, default settings)^2^ and Jalview^20^.

For the non-redundant gene catalog with 147 million genes, LuxI, LuxM, LuxN, HdtS, AHL-acylase and AHL-lactonase were selected based on their phylogenetic clades against reference sequences. The LuxR was screened using the same method as in genome-based analysis, then constructing a phylogenetic tree with reference sequences **(Supplementary Fig. 12)**. The software and parameters used for constructing the phylogenetic tree are the same as those used for the genome-based analysis.

**Identification of quorum sensing genes associated with the DSF system**

For the 3,813 species-level representative MAGs, RpfF sequences were identified using HMMER (v3.3.2) with profiles “PF00378” and “PF16113”^21^, and sequences containing conserved sites Glu114 and Glu164 were selected^22^. Putative RpfC sequences were predicted using the SMART database^23^, retaining those sequences with transmembrane domains, HisKA, HATPase, REC, and HPT domains were retained^24^ **(Supplementary Fig. 13a)**. For RpfG, sequences containing the HD domain and conserved GYP motif were retained using SMART^25^. Conserved sites were visualized using MAFFT (v7.505, default settings)^2^ and Jalview^20^.

For the non-redundant gene catalog with 147 million genes, RpfF and RpfG were screened using the same method as in genome-based analysis **(Supplementary Fig. 13a)**, RpfC and RpfB were selected based on validating their phylogenetic clades against reference sequences **(Supplementary Fig. 13b-c)**. Sequences were aligned using MAFFT (v7.505, default settings)^2^ and trimmed using TrimAL (v1.4.1, default settings)^3^. Maximum-likelihood trees were constructed using FastTree (v2.1.11, default settings)^4^, and all trees were visualized and beautified using iTOL (v6)^5^.

**Identification of quorum sensing genes associated with the c-di-GMP system**

For the 3,813 species-level representative MAGs, DGC sequences were identified using HMMER (v3.3.2) with the profile “PF00990”, and sequences were retained if they contained conserved sites D327, N335, and D344, as well as the inhibitory I-site motif RxxD **(Supplementary Fig. 14)**. PDE sequences were identified using HMMER (v3.3.2) with the profile “PF00563” or by detecting the conserved HD-GYP motif^26^. A phylogenetic tree was constructed to further validate these PDE sequences (**Fig. 4a**). Sequences were aligned using MAFFT (v7.505, default settings)^2^ and trimmed using TrimAL (v1.4.1, default settings)^3^. Maximum-likelihood trees were constructed using FastTree (v2.1.11, default settings)^4^, and all trees were visualized and beautified using iTOL (v6)^5^. Sequences of Clp were predicted using SMART, retaining those sequences with the cAMP-binding domain and conserved residues D70, R166, and D170^27^. Conserved sites were visualized using MAFFT (v7.505, default settings)^2^, Jalview^20^ and WebLogo3 (https://weblogo.threeplusone.com/).

For the non-redundant gene catalog with 147 million genes, sequences of DGC and PDE were screened using the same method as in genome-based analysis, while PDE and Clp were selected based on their phylogenetic clades against reference sequences **(Supplementary Fig. 15)**. The software and parameters used for constructing the phylogenetic tree are the same as those used in the genome-based analysis.

**Identification of quorum sensing genes associated with the PQS system**

For the 3,813 species-level representative MAGs, phylogenetic trees of PhnA, PhnB, PqsE and PqsH were constructed to validate their phylogenetic clades against reference sequences **(Supplementary Fig. 16a-d)**. Sequences were aligned using MAFFT (v7.505, default settings)^2^ and trimmed using TrimAL (v1.4.1, default settings)^3^. Maximum-likelihood trees were constructed using FastTree (v2.1.11, default settings)^4^, and all trees were visualized and beautified using iTOL (v6)^5^. For PqsA, sequences with the SG-TG-PK motif were retained^28^, and PqsD sequences containing Cys112 were selected^29^ **(Supplementary Fig. 16e)**. Conserved sites were visualized using MAFFT (v7.505, default settings)^2^, Jalview^20^ and WebLogo3 (https://weblogo.threeplusone.com/). Deep learning approach AlphaFold2^6^ was applied to predict the three-dimensional structures of PqsL for validation **(Supplementary Fig. 16f)** ^6^. These structures were compared with the structural similarity clustering results obtained from Foldseek (https://cluster.foldseek.com/)^7^, and assessed using Template Modeling Score (TM-score)^8^ and Root Mean Square Deviation (RMSD)^9^. Sequences were considered valid if they met the criteria of TM-score > 0.5^10^ and RMSD < 1. The comparative results were visualized using PyMOL^11^.

For the non-redundant gene catalog with 147 million genes, PqsD and PqsL were screened using the same method as for genome-based analysis, while PhnA, PhnB, PqsA, PqsH and PqsE were selected based on their phylogenetic clades against reference sequences **(Supplementary Fig. 17)**. The software and parameters used for constructing the phylogenetic tree are the same as those used for the genome-based analysis.

**Identification of quorum sensing genes associated with the AHKs system**

For the 3,813 species-level representative MAGs, phylogenetic trees of CqsA and CqsS were constructed to validate their phylogenetic clades against reference sequences **(Supplementary Fig. 18a-b)**. Sequences were aligned using MAFFT (v7.505, default settings)^2^ and trimmed using TrimAL (v1.4.1, default settings)^3^. Maximum-likelihood trees were constructed using FastTree (v2.1.11, default settings)^4^, and all trees were visualized and beautified using iTOL (v6)^5^.

For the non-redundant gene catalog with 147 million genes, phylogenetic trees of CqsA and CqsS were constructed to validate their phylogenetic clades against reference sequences **(Supplementary Fig. 19a-b)**. The software and parameters used for constructing the phylogenetic tree are the same as those used for the genome-based analysis.

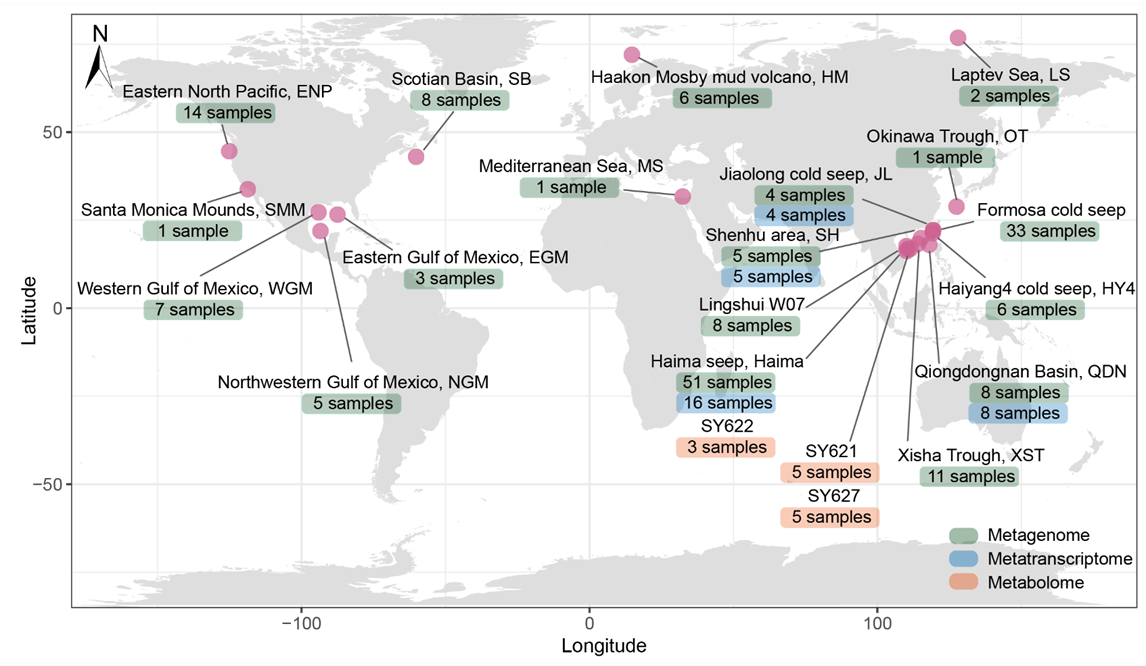

**Supplemental Figure 1. Geographic locations of cold seep sites for metagenomic, metatranscriptomic and metabolomic analyses.** Different colors indicate the type of data collected: green for metagenomic sites, blue for metatranscriptomic sites and orange for metabolomic analysis sites. The world map was generated using the ggplot2 package in R (v4.0.3).

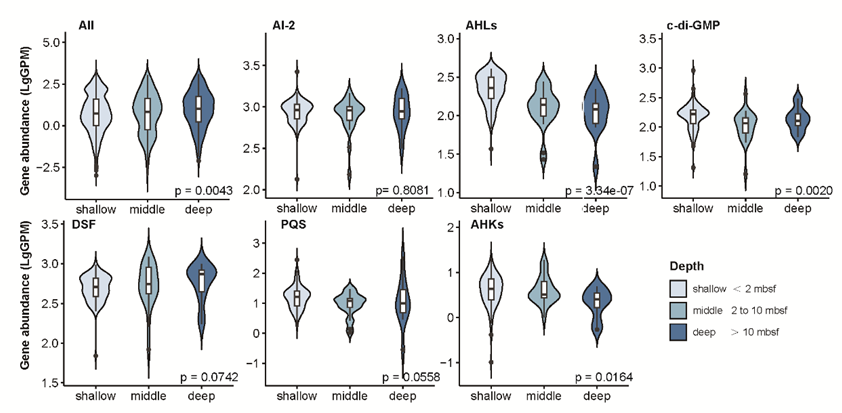

**Supplementary Figure 2. Distribution of the relative abundance of six types of quorum sensing systems across cold seep samples collected from various sediment depth layers.** Sediment depths are categorized into three groups: surface (< 2 meters below seafloor, mbsf), shallow (2-10 mbsf), and deep (> 10 mbsf). The vertical axis represents the lg-transformed values of gene relative abundance (lg-transformed genes per million, lgGPM) at different sediment layer depths. Kruskal-Wallis rank-sum tests were used to determine differences in the abundance of **(a)** all quorum sensing systems and **(b-g)** individual types of quorum sensing systems at different sediment layer depths, with *P* values calculated to determine the statistical significance of the observed variations. Violin plots show: the center line representing the median, box limits indicating the 25th and 75th percentiles, and whiskers extending 1.5 times the interquartile range from the 25th and 75th percentiles. Detailed statistical data are available in **Supplementary Table 7.**

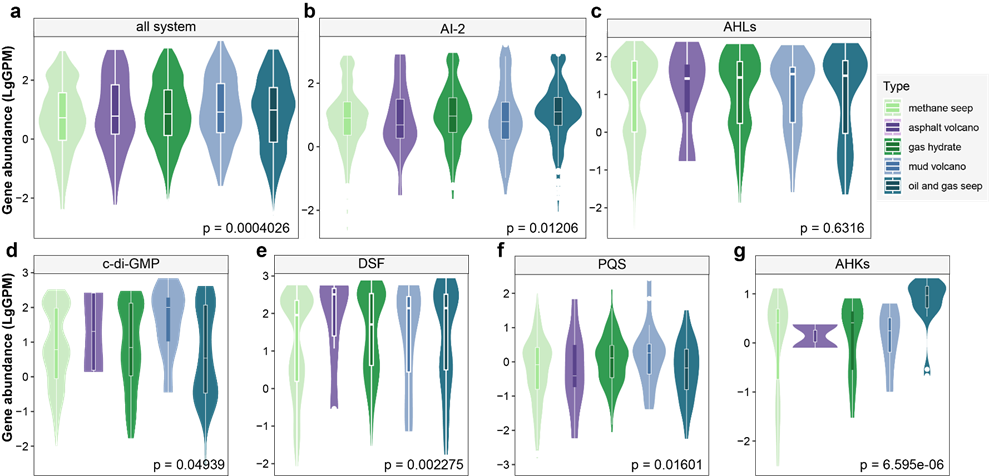

**Supplementary Figure 3. Distribution of relative abundance of six quorum sensing systems across 165 cold seep sediment samples collected from five cold seep types.** Kruskal-Wallis rank-sum tests were used to determine differences in the abundance of **(a)** all quorum sensing systems and **(b-g)** individual types of quorum sensing systems across different cold seep types. *P* values indicate the significance of these differences. Violin plots show: the center line representing the median, box limits indicating the 25th and 75th percentiles, and whiskers extending 1.5 times the interquartile range from the 25th and 75th percentiles. The vertical axis represents the log-transformed values of gene relative abundance (lg-transformed genes per million, lgGPM).

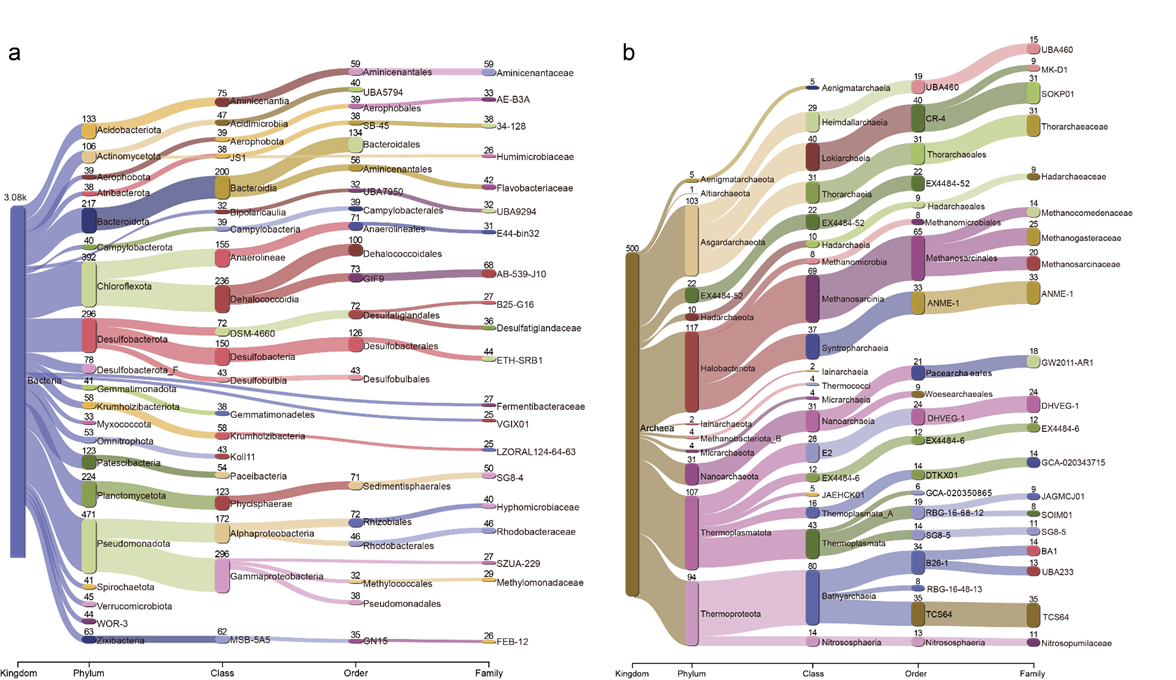

**Supplementary Figure 4. Sankey plots showing cold seep microorganisms with quorum sensing genes across different taxonomic levels. (a)** Archaeal MAGs and **(b)** bacterial MAGs. Numbers indicate the quantity of MAGs recovered for each lineage. Detailed classification information for the MAGs is provided in **Supplementary Table 8**.

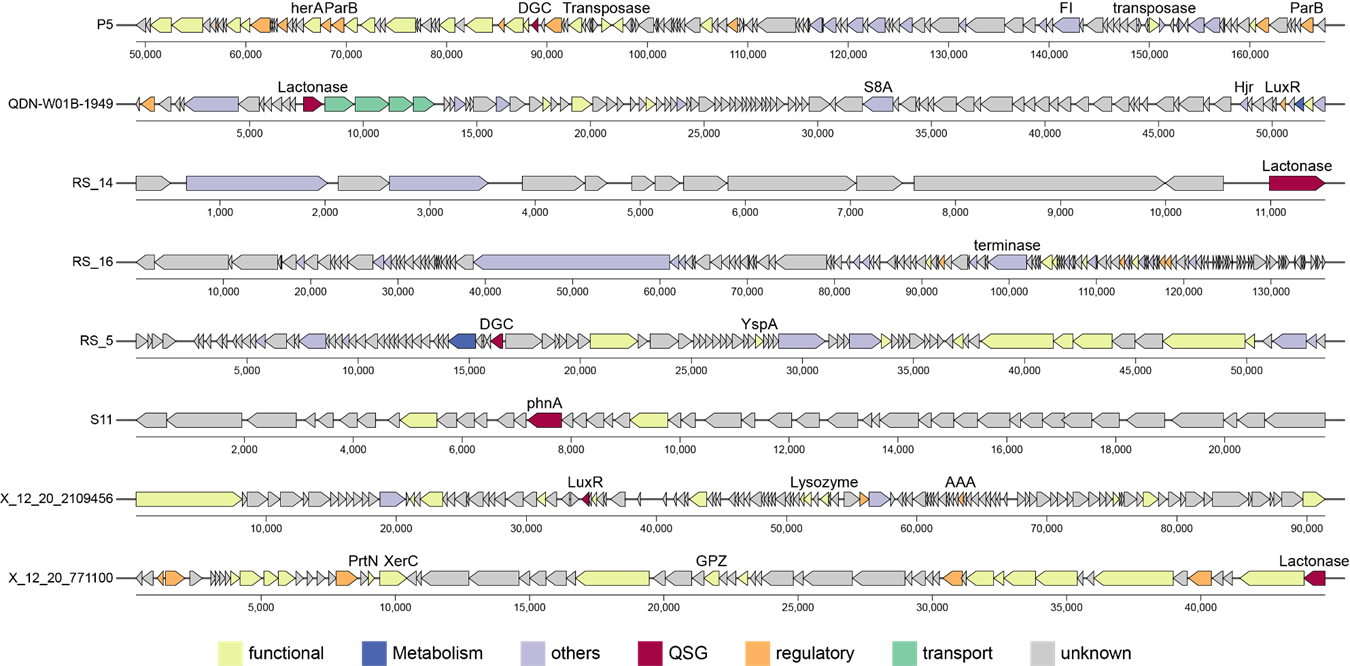

**Supplementary Figure 5. Genome synteny of eight representative viral contigs containing quorum sensing genes and their upstream and downstream genes.** Different colors represent various gene types: functional genes (yellow-green), metabolism-related genes (blue), other annotated genes (purple), quorum sensing genes (red), regulatory genes (yellow), transport-related genes (green), and unknown genes (gray).

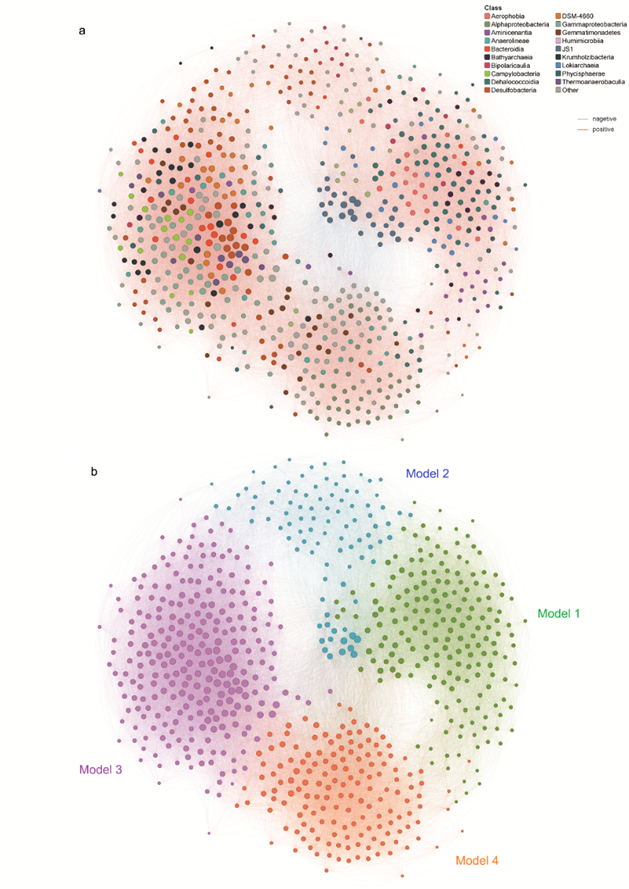

**Supplementary Figure 6. Co-occurrence networks for microorganisms significantly correlated with quorum sensing genes.** Node size reflects the degree of connectivity. Nodes are colored by class in **(a)** and by model in **(b).** In panel **(a)**, red edges indicate significant positive correlations, while blue edges represent significant negative correlations.

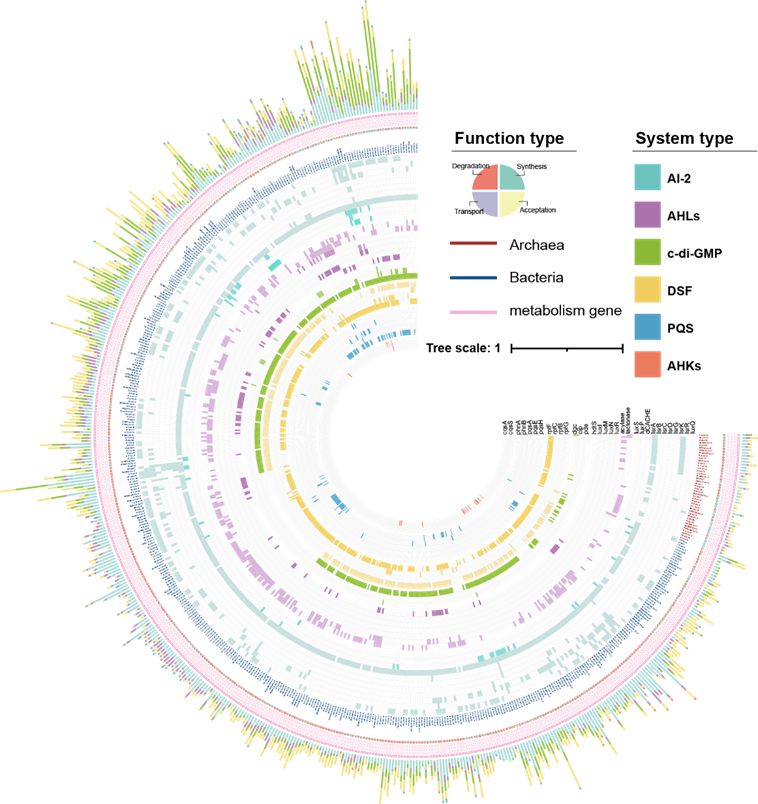

**Supplementary Figure 7.** **Distribution of quorum sensing related genes in organohalide reducers.** The innermost layer with colored squares indicates the presence of quorum sensing genes, while different colors representing different types of quorum sensing systems. Archaea and bacteria are distinguished by label colors, with red labels indicating archaea and blue labels indicating bacteria. Pie charts display the proportion of four functional gene categories. The next outer layer with pink indicates the presence of key metabolic marker genes. The outermost layer presents a stacked bar chart showing the proportion of six quorum sensing systems in metabolic microbial lineages.

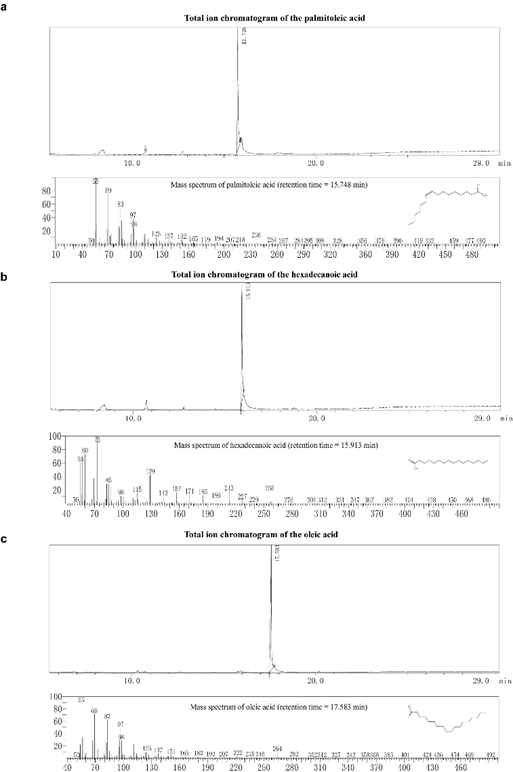

**Supplementary Figure 8. Mass chromatograms of hexadecanoic acid, palmitoleic acid, and oleic acid standards analyzed using GC-MS. (a)** Top: total ion chromatogram (TIC) of hexadecanoic acid standard, with a retention time of 15.915 min; Bottom: mass spectrum of hexadecanoic acid standard at the chromatographic signal corresponding to a retention time of 15.915 min. **(b)** Top: TIC of palmitoleic acid standard, with a retention time of 15.750 min; Bottom: mass spectrum of palmitoleic acid standard at the chromatographic signal corresponding to a retention time of 15.750 min. **(c)** Top: TIC of oleic acid standard, with a retention time of 17.580 min; Bottom: mass spectrum of oleic acid standard at the chromatographic signal corresponding to a retention time of 17.580 min.

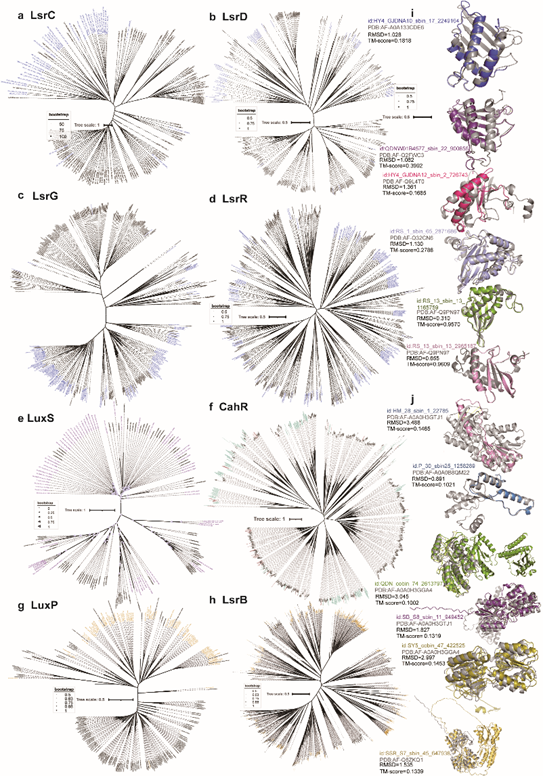

**Supplementary Figure 9. Quorum sensing genes related to AI-2 system identified from cold seep microbiomes. (a-h)** Maximum-likelihood phylogenetic trees of LsrC, LsrD, LsrG, LsrR, LuxS, CahR, LuxP and LsrB. Colored labels represent target genes from cold seep microbiomes, while black labels indicate reference genes. The scale bar represents the mean number of substitutions per site. The three-dimensional structures of LuxS **(i)** and LuxP **(j)** were aligned with the structures of their corresponding reference proteins for comparison. Colored are structures of target sequences that cluster poorly with reference genes in the phylogenetic tree, gray are the structures of reference proteins. TM-score and RMSD values were used to quantify the degree of structural similarity between the target and reference proteins.

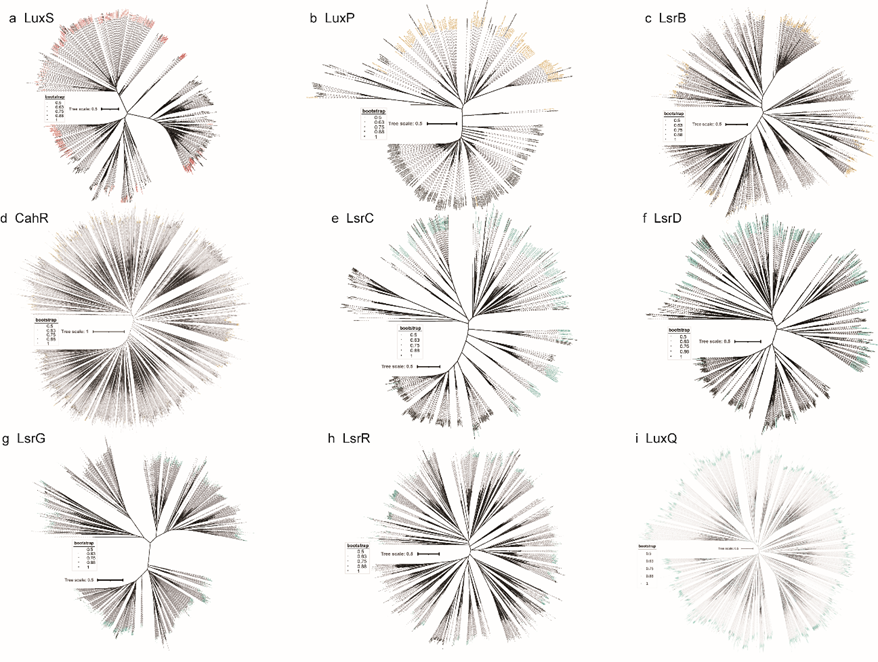

**Supplementary Figure 10. Maximum-likelihood phylogenetic trees of quorum sensing genes related to AI-2 system identified from the non-redundant gene catalog. (a-i)** Phylogenetic trees of LuxS, LuxP, LsrB, CahR, LsrC, LsrD, LsrG, LsrR and LuxQ. Colored labels represent target genes, while black labels indicate reference genes. The scale bar denotes the mean number of substitutions per site.

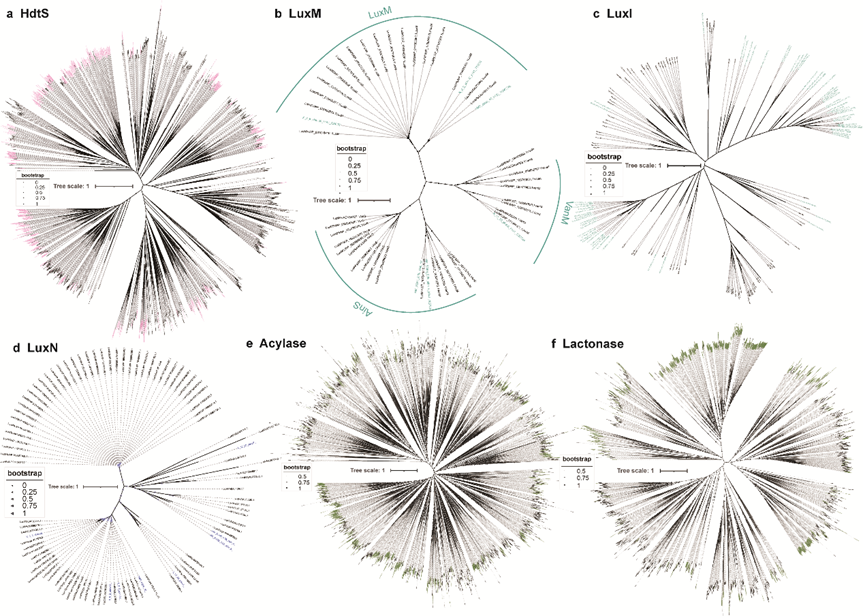

**Supplementary Figure 11. Maximum-likelihood phylogenetic trees for quorum sensing genes related to AHLs system identified from cold seep microbiomes. (a-f)** Phylogenetic trees of HdtS, LuxM, LuxI, LuxN, AHL-acylase and AHL-lactonase. Colored labels represent target genes from cold seep microbiomes, while black labels indicate reference genes. The scale bar indicates the mean number of substitutions per site.

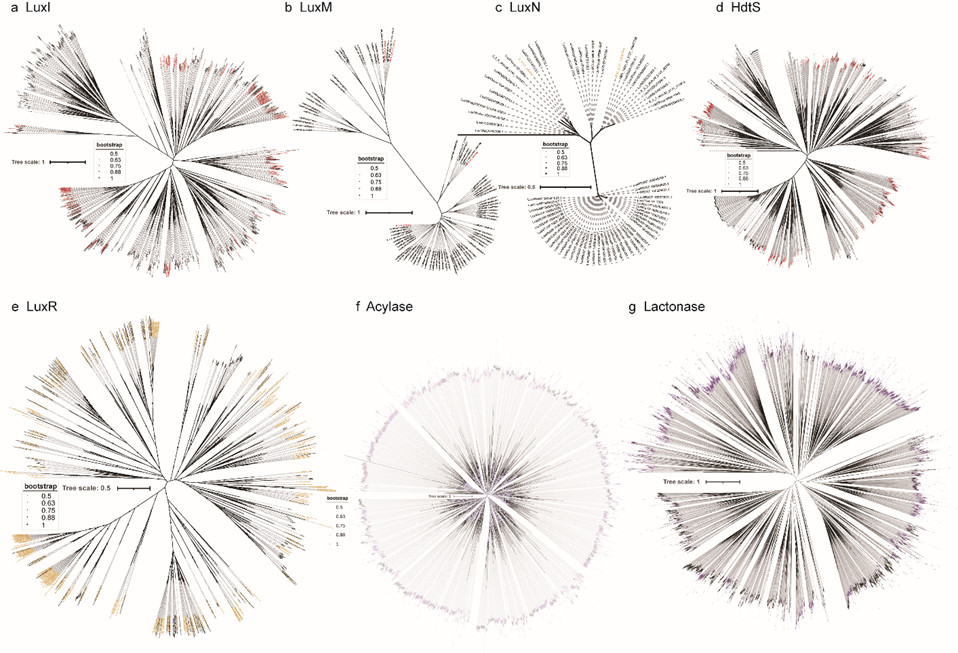

**Supplementary Figure 12. Maximum-likelihood phylogenetic trees for quorum sensing genes related to AHLs system identified from the non-redundant gene catalog. (a-g)** Maximum-likelihood phylogenetic trees for LuxI, LuxM, LuxN, HdtS, LuxR, AHL-acylase and AHL-lactonase. Colored labels represent target genes, while black labels indicate reference genes. The scale bar denotes the mean number of substitutions per site.

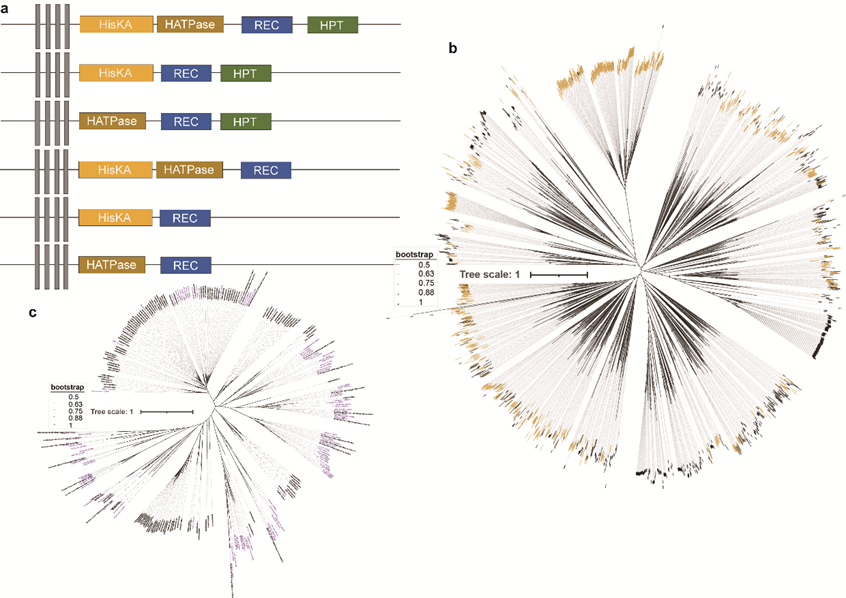

**Supplementary Figure 13. Quorum sensing genes related to DSF system identified from the non-redundant gene catalog. (a)** Gene structure features of *rpfC* are illustrated, showing domain arrangements such as HisKA, HATPase, REC, and HPT. **(b-c)** Maximum-likelihood phylogenetic trees for RpfC and RpfB. Colored labels represent target genes, while black labels indicate reference genes. The scale bar indicates the mean number of substitutions per site.

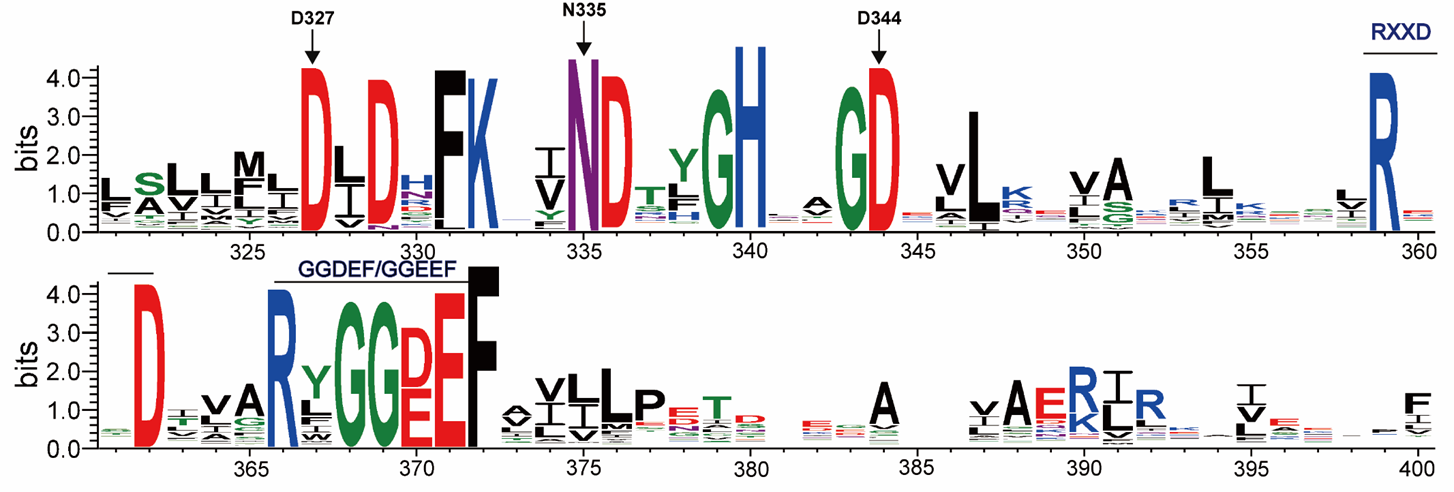

**Supplementary Figure 14. The DGC domain conservation model related to c-di-GMP system identified from cold seep microbiomes.** Multiple sequence alignment logo of 8,367 DGC domain sequences selected from cold seep microbiomes, highlighting 10 conserved sites of the DGC gene domain, including D327, N335, D344, RxxD, and GGDEF/GGEEF motifs. The multiple sequence alignment was performed using MAFFT and visualized with WebLogo3.

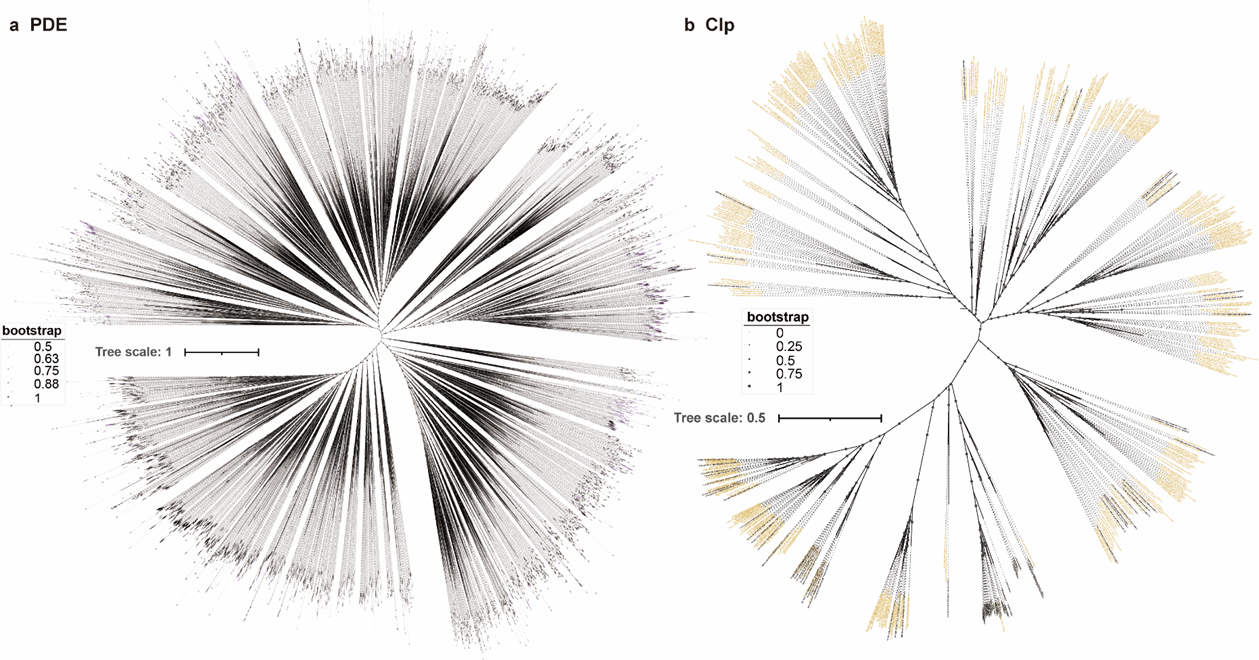

**Supplementary Figure 15. Maximum-likelihood phylogenetic trees for PDE and Clp related to c-di-GMP system identified from the non-redundant gene catalog.** Phylogenetic trees are shown for PDE **(a)** and Clp **(b).** Colored labels represent target genes, while black labels indicate reference genes. The scale bar shows the average number of substitutions per site.

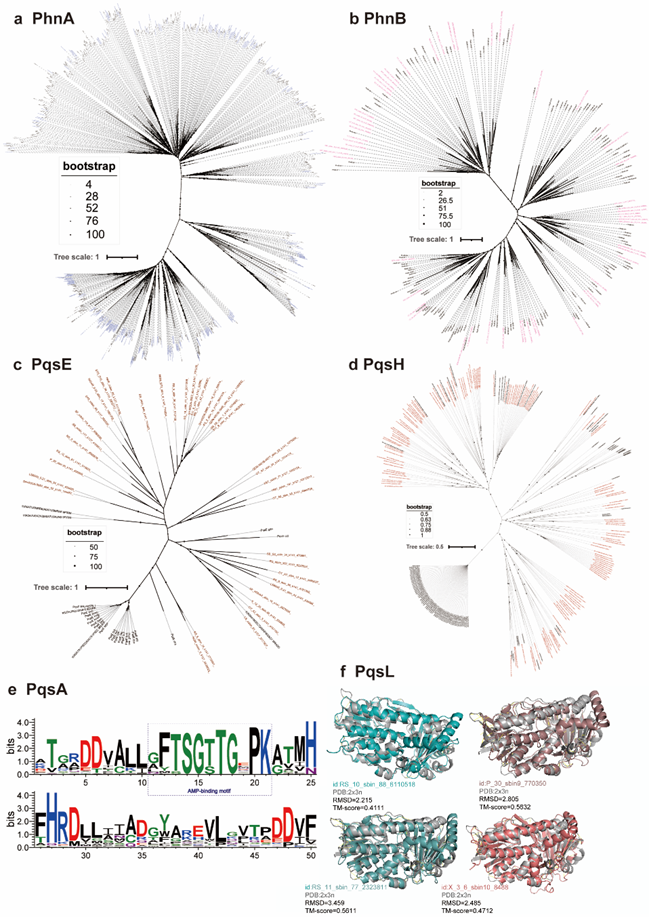

**Supplementary Figure 16. Quorum sensing genes related to PQS system identified from cold seep microbiomes. (a-d)** Maximum-likelihood phylogenetic trees for PhnA, PhnB, PqsE and PqsH. Colored labels represent target genes from cold seep microbiomes, while black labels indicate reference genes. The scale bar represents the mean number of substitutions per site. **(e)** The PqsA domain conservation model, showing key conserved sites. **(f)** Structural alignments of PqsL with their corresponding reference proteins. The alignment results include TM-score and RMSD values, which quantify the degree of structural similarity. Colored structures represent target protein structures, while gray structures represent reference protein structures.

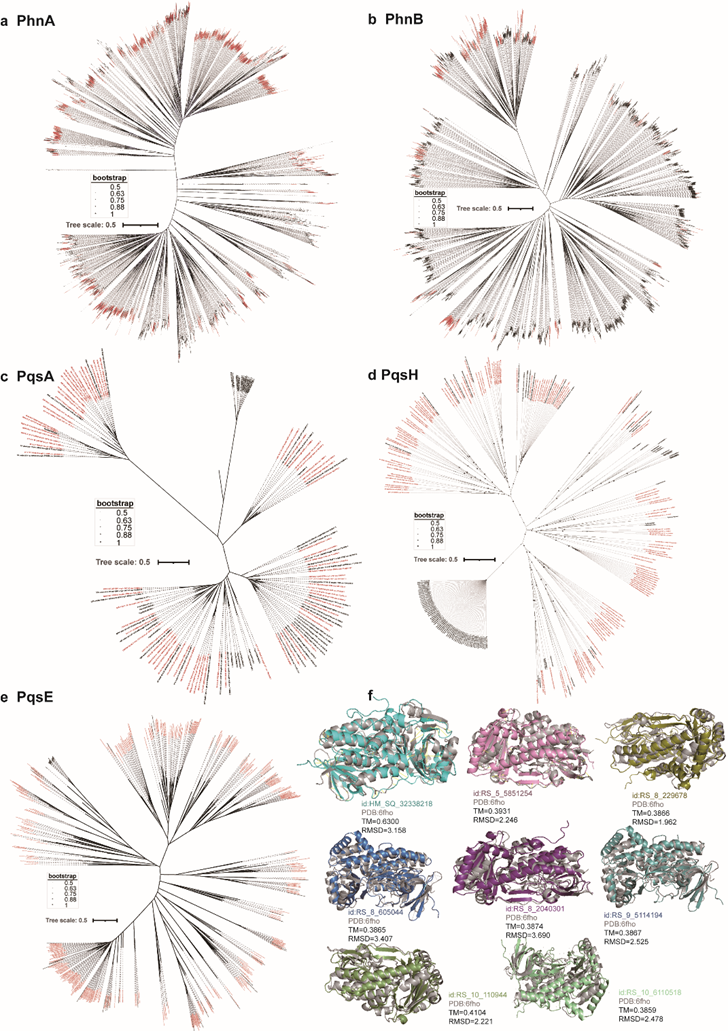

**Supplementary Figure 17.** **Quorum sensing genes related to PQS system identified from the non-redundant gene catalog. (a-e)** Maximum-likelihood phylogenetic trees for PhnA, PhnB, PqsA, PqsH and PqsE. Colored labels represent target genes, while black labels indicate reference genes. **(f)** Structural alignments of PqsL with their corresponding reference proteins. TM-score and RMSD values were used to quantify the degree of structural similarity.

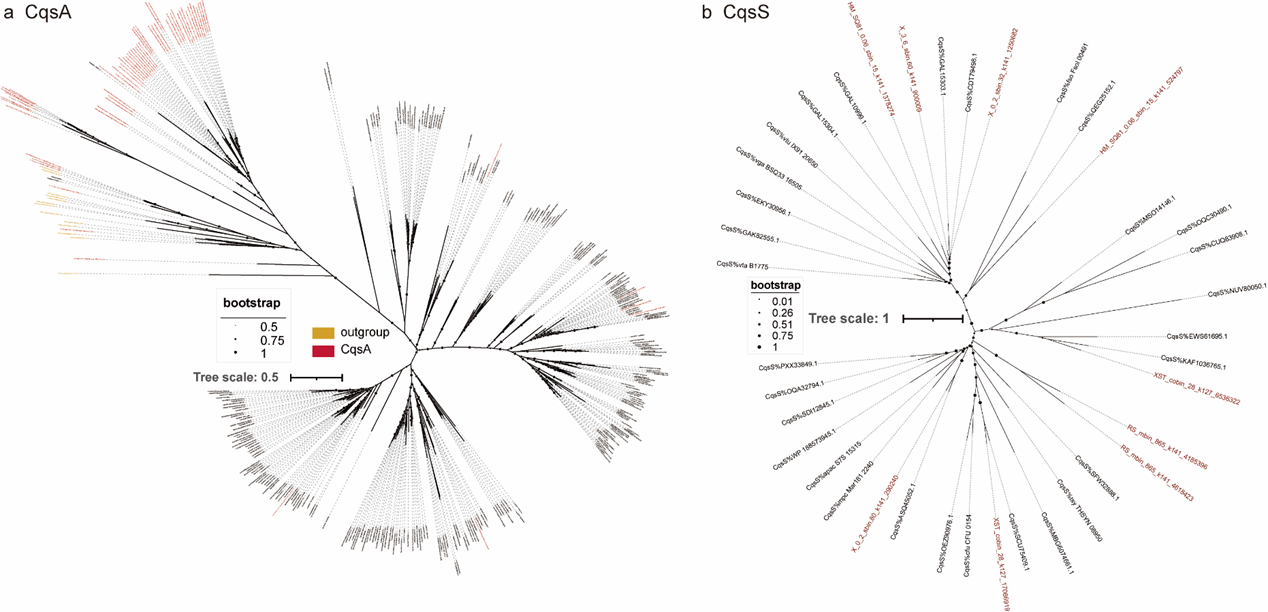

**Supplementary Figure 18. Maximum-likelihood phylogenetic trees for quorum sensing genes related to the AHKs system identified from cold seep microbiomes.** Phylogenetic trees are shown for CqsA **(a)** and CqsS **(b).** Colored labels represent target genes, while black labels indicate reference genes. The scale bar indicates the mean number of substitutions per site.

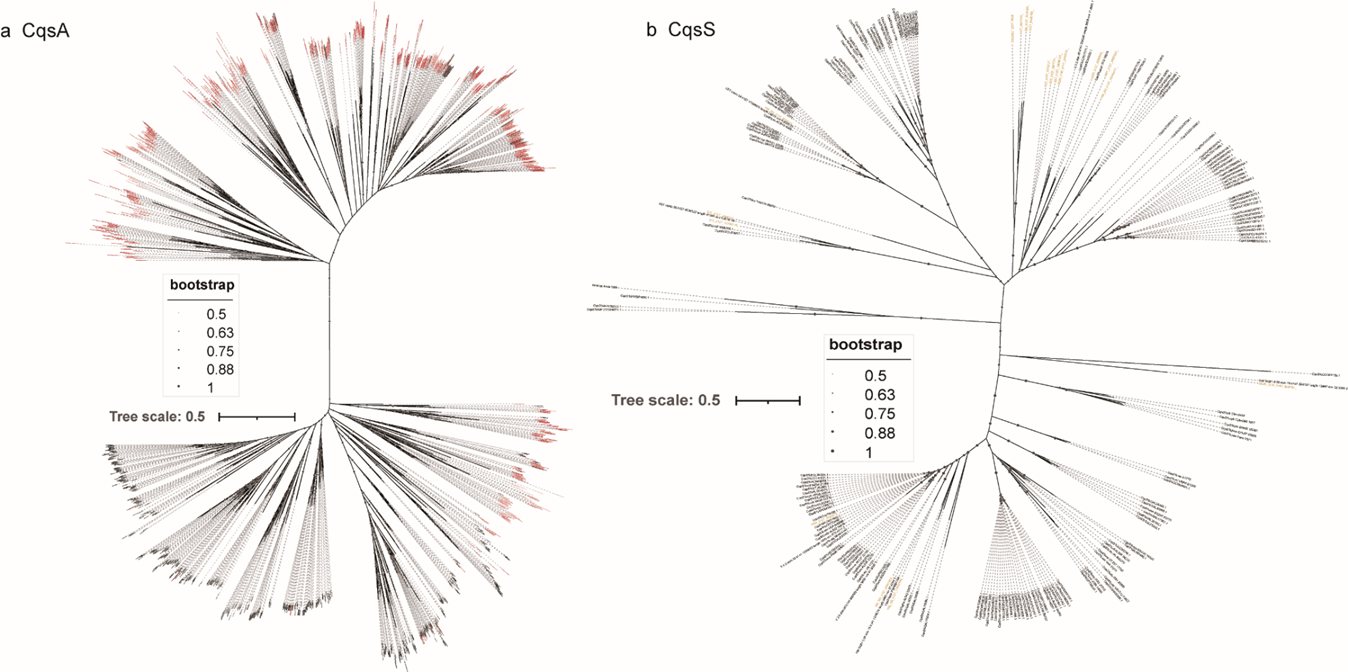

**Supplementary Figure 19. Maximum-likelihood phylogenetic trees for quorum sensing genes related to AHKs system identified from the non-redundant gene catalog.** Phylogenetic trees are shown for CqsA **(a)** and CqsS **(b)**. Colored labels represent target genes, while black labels indicate reference genes. The scale bar indicates the mean number of substitutions per site.

**Table S1.** Summary of quorum sensing protein screening methods.

**Table S2.** Detailed information on 299,355 quorum sensing proteins in the non-redundant gene catalog.

**Table S3.** Average abundance of quorum sensing systems.

**Table S4.** Average abundance of quorum sensing proteins.

**Table S5.** Pairwise significance tests for relative abundance among six quorum sensing systems using the Wilcoxon test.

**Table S6.** Pairwise significance tests for relative abundance among quorum sensing proteins using the Wilcoxon test.

**Table S7.** Pairwise significance tests for the relative abundance of quorum sensing proteins across three depth layers using the Wilcoxon test.

**Table S8.** Detailed information on 32,500 quorum sensing proteins in 3,576 cold seep metagenome-assembled genomes (MAGs).

**Table S9.** Comprehensive details of 84 quorum sensing proteins identified in viral genomes using HMMER from the QSP database.

**Table S10.** Pairwise correlation analysis of quorum sensing proteins based on Spearman’s rank correlation analysis.

**Table S11.** Network characteristics of quorum sensing systems.

**Table S12.** Quorum sensing proteins and metagenome-assembled genomes (MAGs) with significant correlations.

**Table S13.** Nodes of metagenome-assembled genomes (MAGs) with higher degrees (8-14) and prediction of hub nodes in microbial co-occurrence networks.

**Table S14**. Statistical data for linear regression between the abundance of *dsrA*, *mcrA*, *nifH*, and *rdhA* and quorum sensing genes.

**Table S15.** Distribution of quorum sensing proteins in functionally key microbial groups and taxonomic classification of metagenome-assembled genomes (MAGs).

**Table S16.** Diversity of dCACHE_1-containing AI-2 receptors predicted using the Pfam database.

**Table S17.** List of identified *lux* box sequences and their downstream genes predicted in functionally key microbial groups using FIMO-MEME.
